# The Role of Smooth Muscle Cell Heterogeneity in Cerebral Autoregulation: A Multi-Scale Physics-Based Modeling Study

**DOI:** 10.64898/2026.09.14.751526

**Authors:** Nele Demeersseman, Lauranne Maes, Bart Depreitere, Nele Famaey

## Abstract

**Background:** Cerebral autoregulation stabilizes cerebral blood flow over a range of cerebral perfusion pressures, but the precise shape of the pressure-flow relationship remains debated. The classical triphasic pressure-flow relationship was recently challenged by experiments demonstrating a quadriphasic response, hypothesized to arise from vessel-size-dependent pressure-diameter responses. We tested this hypothesis and investigated whether these size-dependent responses originate from heterogeneity in smooth muscle cell (SMC) abundance, SMC behavior, or neither.

**Methods:** We developed a computational multi-scale physics-based model of cerebral autoregulation linking SMC activity to vessel-scale diameter regulation and organ-scale blood flow. Four scenarios were evaluated: passive vessels, homogeneous SMC abundance and behavior, heterogeneous SMC abundance, and heterogeneous SMC behavior. Predicted pressure-diameter responses and pressure-flow relationships were compared across scenarios and against experimental observations.

**Results:** In contrast to passive vessels, homogeneous SMC activation produced partial flow stabilization, highlighting the key role of SMCs in autoregulation. However, only heterogeneous SMC behavior reproduced the experimentally observed vessel-size-dependent trends in pressure-diameter responses. This scenario also showed the best agreement with the experimental organ-scale pressure-flow relationship (R^2^ = 0.93, nRMSE = 5.96%).

**Conclusion:** The model suggests that vessel-size-dependent SMC behavior underlies vessel-size-dependent pressure-diameter responses and shapes the relationship between cerebral perfusion pressure and cerebral blood flow.

## 1. Introduction

The brain is highly sensitive to fluctuations in blood flow, as even brief disruptions in oxygen and nutrient delivery can impair neuronal function and lead to tissue injury. To safeguard cerebral homeostasis, the cerebral circulation actively regulates cerebral blood flow (CBF) through a set of tightly coordinated mechanisms, together known as CBF regulation. These mechanisms adjust cerebrovascular resistance in response to changes in cerebral perfusion pressure, arterial blood gasses, and other physiological stimuli, thereby stabilizing CBF across a wide range of systemic conditions^1,2^. The specific mechanisms ensuring relatively stable CBF under variations in cerebral perfusion pressure are collectively referred to as cerebral autoregulation. Central to autoregulation is the myogenic response, whereby cerebrovascular smooth muscle cells constrict or dilate in response to changes in vascular pressure^1^.

At the macroscopic level, cerebral autoregulation is traditionally described using the pressure-flow curve introduced by Lassen in 1959^3^. In this three-phasic description, CBF remains relatively constant over a broad range of cerebral perfusion pressures (50 to 150 mmHg) and becomes linearly pressure dependent outside this range. While this description has been widely influential, accumulating evidence suggests that it does not fully capture the complexity of cerebral autoregulation^2,4,5^. In particular, a recent experimental study in piglets by Klein et al.^5^ reported a quadriphasic pressure-flow relationship, characterized by a more narrow plateau of near-constant flow (39 to 70 mmHg) followed by a gradual and subsequently steeper increase in CBF as myogenic control was lost. In addition, a recent review by Wang et al.^2^ concluded that the mechanisms underlying cerebral autoregulation remain incompletely understood, limiting our ability to interpret both physiological vascular control and its impairment in pathological conditions such as dementia, stroke, and traumatic brain injury^2,6,7^.

A key step toward understanding the complexity of autoregulation is to determine how pressure-dependent responses vary across vessel sizes within the cerebrovascular network. Indeed, according to Klein et al.^5^, vessel-size-dependent pressure-diameter responses may underlie the observed quadriphasic pressure-flow relationship. They further hypothesized that these size-dependent responses stem from differences in smooth muscle cell (SMC) content along the vascular tree but were unable to verify this. An alternative, and not mutually exclusive, explanation is that vessel-size-dependent pressure responses reflect functional differences in the behavior of SMCs across vessel sizes. Supporting this view, Cipolla et al.^8^ demonstrated distinct ion channel activity in SMCs from cerebral arterioles compared to cerebral arteries in rats. While these findings provide evidence for intrinsic cellular heterogeneity, their impact on organ-scale pressure-flow relationships remains unexplored.

Experimentally studying the effects of SMC heterogeneity (in abundance and/or function) on cerebral autoregulation is challenging because the underlying mechanisms span multiple organizational scales. Cerebral autoregulation is typically assessed at the level of global CBF using *in vivo* approaches^5,9^, while mechanistic insights into SMC function are derived primarily from isolated vessels or cellular preparations^8,10^. This scale separation complicates direct translation from cellular observations to organ-scale pressure-flow behavior, highlighting the need for integrative approaches that can explicitly link SMC properties to vascular network hemodynamics within a unified framework.

Computational modeling addresses this need by providing a controlled framework in which cellular mechanisms, vessel wall mechanics, and network-level hemodynamics can be systematically integrated and analyzed^11–13^. While modeling approaches have already proven useful in other vascular fields^14,15^, their application to cerebral autoregulation remains limited. Many existing vascular models neglect SMC dynamics entirely or represent them in a simplified manner^16^, limiting their transferability to autoregulation where active SMC control plays a central role^1^. Models specifically developed for cerebral autoregulation, on the other hand, are often compartmental, reducing the vascular network to a small number of lumped elements, precluding investigation of vessel-size-dependent responses^7,11,17^. Network-based models can overcome this limitation but the few that currently exist typically rely on phenomenological descriptions of the vessel wall and/or cellular processes^7,12^, restricting mechanistic interpretation of SMC contributions to autoregulation.

Motivated by these gaps, we developed a computational multi-scale physics-based model of cerebral autoregulation that explicitly links processes across cellular, vessel, and organ scales. The model integrates existing experimental and theoretical knowledge to enable systematic investigation of how local myogenic responses shape global pressure-flow behavior. Using this framework, we examine whether vessel-size-dependent pressure responses can give rise to the reported quadriphasic autoregulatory curve, as suggested by Klein et al.^5^, and whether these size-dependent responses emerge from structural heterogeneity in SMC abundance, functional heterogeneity in SMC behavior, or neither.

## 2. Materials and methods

### 2.1. Model structure and governing equations

#### 2.1.1. Model overview

The model is formulated as a multi-scale representation of the cerebral vasculature, spanning organ, vessel, microstructural, and molecular scales (Figure 1). It is designed to study static cerebral autoregulation by computing steady-state pressure-flow relationships under slow changes in cerebral perfusion pressure (CPP).

**Figure 1.**
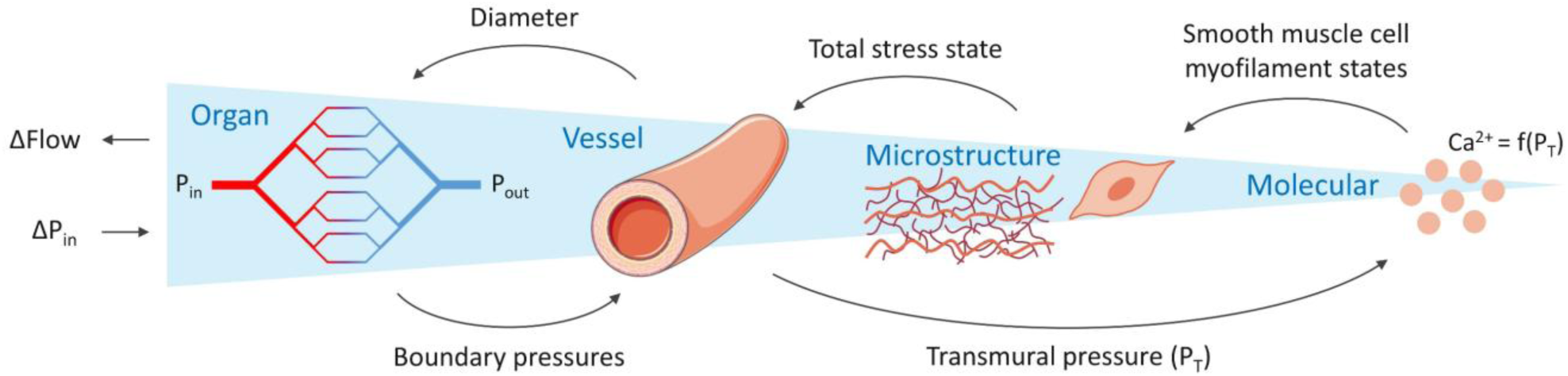
Schematic overview of the multi scale cerebral autoregulation model, illustrating the coupling between organ, vessel, microstructural, and molecular scales. Some of the icons used in this schematic were obtained from Servier Medical Art (https://smart.servier.com), licensed under CC BY 4.0 (https://creativecommons.org/licenses/by/4.0/).

As illustrated in Figure 1, CPP is varied through stepwise changes in inlet pressure at the organ scale. Each change induces a redistribution of pressures across the vascular network, thereby defining the mechanical load on individual vessels. This loading is propagated to the molecular scale, where pressure-dependent signaling pathways regulate SMC myofilament activation. At the microstructural scale, this activation generates active SMC stress, which combines with passive elastin and collagen stresses to define the mechanical state of the vessel wall. By solving for mechanical equilibrium, the corresponding vessel diameters are obtained and returned to the organ-scale network, where they lead to a redistribution of cerebrovascular resistance and pressures. This coupled multi-scale process is iterated until convergence of pressure and diameter distributions is achieved. The convergence criteria can be found in Supplementary Material S1.4. Once converged, cerebral blood flow is computed. All computations were done in MATLAB R2024b (The Mathworks Inc., Natick, Massachusetts, USA).

#### 2.1.2. Organ scale

The organ-scale vascular network consists of an arterial tree, a capillary bed, and a venous tree^7^. The arterial tree starts at the middle cerebral artery, with a prescribed diameter *D_in_*, and continues through symmetric bifurcation^18^. Daughter vessel diameters *D_d_* are computed from the parent diameter *D_p_* using Murray’s law (Equation (1)^19^ with a diameter-dependent exponent *n_M_*(*D*) (Equation 2) derived from reference small- and large-vessel diameter-exponent pairs ((*D_M_*_,*s*_, *n_M_*_,*s*_) and (*D_M_*_,*l*_, *n_M_*_,*l*_))^20,21^. Branching continues until a terminal arteriole diameter is reached, defined as a fixed multiple (*λ_a_*_/*c*_) of the capillary diameter (*D_c_*)^12,22,23^. Consistent with the data of Cassot et al.^21^, the reference small-vessel diameter *D_M_*_,*s*_ corresponds to a terminal arteriole (*D_M_*_,*s*_ = *λ_a_*_/*c*_*D_c_*).

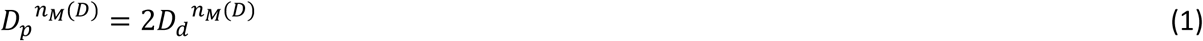

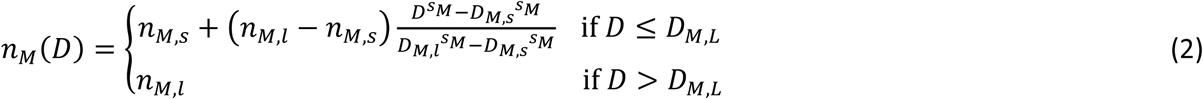

The venous tree is constructed analogously to the arterial tree^21^, starting from a venous diameter defined as a multiple (*λ_v_*_/*a*_) of the arterial inlet diameter to account for the larger caliber of veins^22,24^. Venous branching is continued until the number of terminal venules matches the number of terminal arterioles. Arterial and venous trees are connected through a capillary bed consisting of *N_c_* parallel, non-interconnecting vessels per terminal arteriole. Vessel lengths and wall thicknesses are assigned using length-to-diameter (*λ_L_*_/*D*_) and thickness-to-diameter ratios (*λ_t_*_/*D*_) specific to arterial, capillary, and venous segments^22,25^.

Hydraulic resistance is computed assuming laminar flow, with apparent blood viscosity described using the *in vivo* diameter-dependent relations of Pries et al.^26^. Boundary conditions are defined by mean inlet (*P_in_*), outlet (*P_out_*), and intracranial (*P_ICP_*) pressures. Static cerebral autoregulation is assessed by applying stepwise 10% changes in CPP relative to baseline. These are translated directly into variations in *P_in_* as *P_ICP_* and the difference between MAP and *P_in_* are both assumed constant (CPP = MAP - *P_ICP_*). The venous outlet pressure *P_out_* is also assumed constant throughout the simulations. Steady-state pressures at the vascular nodes are obtained by enforcing conservation of flow at each node. Blood flow is computed using Hagen-Poiseuille. Governing equations are provided in Supplementary Material S1.1.

#### 2.1.3. Vessel scale

For each vessel, proximal and distal luminal pressures obtained from the organ-scale network are averaged to define the network-prescribed luminal pressure *P_l_*_,*network*_. To determine the vessel radius that can mechanically sustain this pressure, the vessel wall is modeled as a pressurized thick-walled cylinder such that the mechanical equilibrium in the radial direction yields^13,15^

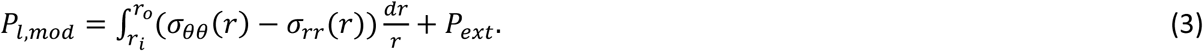

Here, *P_l_*_,*mod*_ is the luminal pressure according to the model, *r_i_* and *r_o_* are the inner and outer radii of the vessel in its current configuration, *P_ext_* is the intracranial pressure, and *σ_θθ_* and *σ_rr_* denote the circumferential and radial Cauchy stresses^15,27^. These stresses depend on the vessel radius, which is obtained by solving for *P_l_*_,*mod*_ = *P_l_*_,*network*_ using a numerical root-finding method. The corresponding vessel thickness follows from the assumptions of incompressibility and constant vessel length^28^.

The exact relationship between wall stress and vessel radius depends on vessel type. Capillary and venous segments are treated as non-regulating vessels^7^ and described using a lumped passive wall formulation. Arterial segments are treated as regulating vessels^7^, where total wall stress (**σ***_tot_*) is defined as the volume-fraction-weighted sum of the stress contributions carried by elastin, collagen, and smooth muscle cells^29^:

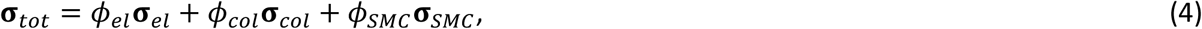

with *φ_i_* the volume fractions. The constitutive descriptions of the different stresses are given in the following section.

#### 2.1.4. Microstructural scale

For constituent *i*, the Cauchy stress (**σ***_i_*) under incompressible hyperelasticity is given by

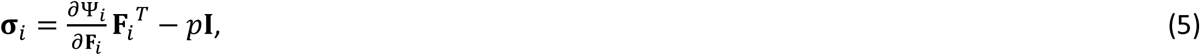

where **I** is the identity tensor and *p* is a Lagrange multiplier enforcing incompressibility^15^. **Ψ***_i_* is the strain-energy density function (SEDF) that characterizes the mechanical response of wall constituent *i* as a function of its deformation measure **F***_i_* (see Supplementary Material S1.2. for details)^29^. Elastin is modeled using a Neo-Hookean SEDF,

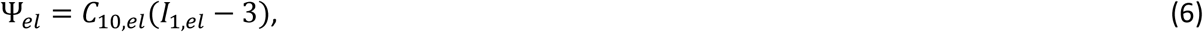

where *C*_10,*el*_ represents elastin stiffness and *I*_1,*el*_ is the first invariant derived from **F***_el_*^29^.

Collagen is modeled as an anisotropic fiber-reinforced material consisting of two symmetric fiber families. For each fiber family *j*, the SEDF is given by

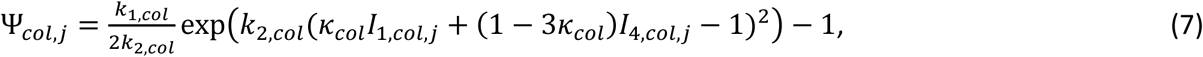

where *k*_1,*col*_ and *k*_2,*col*_ control fiber stiffness and nonlinearity, *κ_col_* describes fiber dispersion, *I*_1,*col*,*j*_ is the first invariant derived from **F***_col_*_,*j*_, and *I*_4,*col*,*j*_ represents the squared stretch along the average fiber direction^29^.

Similar to collagen, the SMCs are represented by two symmetric families. The SEDF of an individual SMC family is given by

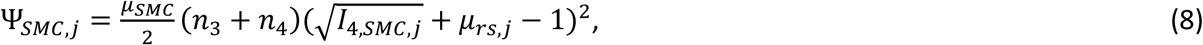

where *μ_SMC_* is an active stiffness parameter, *I*_4,*SMC*,*j*_ is the invariant associated with the SMC contractile direction, and *μ_rs_*_,*j*_ defines the normalized relative sliding between SMC filaments^29,30^. The parameters *n*_3_ and *n*_4_ reflect the fraction of SMC filaments in active states and are obtained from the molecular scale model (Section 2.1.5). In the calibrated model, the SMC family angles are set to 0^∘^ (Section 2.3), such that both families actually coincide. The influence of non-zero SMC family angles is evaluated in the sensitivity analysis.

Capillary and venous segments are described using Neo-Hookean SEDFs (as in Equation 6) with vessel-type-specific material parameters *C*_10,*c*_ and *C*_10,*v*_, respectively.

#### 2.1.5. Molecular scale

Active SMC contraction is described using the Hai-Murphy model^31^, in which myosin cross-bridges transition between four biochemical states (*n*_1_-*n*_4_) based on rate constants *k*_1_-*k*_7_. The full system of equations is provided in Supplementary Material S1.3. Building on this framework, we introduce a dependency between the rate of phosphorylation *k*_1_ (and *k*_6_=*k*_1_) and the intracellular calcium concentration ([*Ca*^2+^]) in a two-stepped procedure. First, the phosphorylation rate is directly dependent on calmodulin (*CaM*) activation (i.e. the binding of *CaM* with *Ca*^2+^ to form *CaCaM*) as

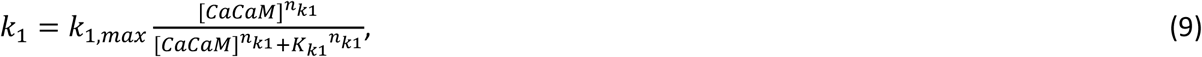

with *k*_1,*max*_ the maximum rate, *K_k_*_1_ the half-activation constant, and *n_k_*_1_ a Hill exponent^29,30^. Secondly, in contrast to Maes et al.^29^ and Murtada et al.^30^, which assumed a linear relation between [*Ca*^2+^] and [*CaCaM*], calmodulin activation is described using a Hill function to better reflect the cooperative binding of calcium to calmodulin^32,33^:

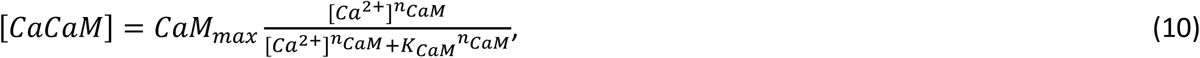

with *CaM_max_* the maximal calcium-calmodulin concentration, *K_CaM_* the half-activation constant, and *n_CaM_* a Hill exponent.

Intracellular calcium itself varies with transmural pressure *P_T_* = *P_l_*_,*network*_ − *P_ext_*, thereby closing the multi-scale loop^13,34^. Their relationship can be represented with varying degrees of detail, depending on the desired balance between model simplicity and physiological fidelity. In its simplest form, a single pressure-calcium curve can be applied uniformly across all vessels, assuming homogeneous SMC behavior. Greater physiological fidelity can be achieved by incorporating vessel-size-dependent pressure-calcium responses, consistent with experimental observations^8,10^. The formulations investigated in this study are introduced in the simulation scenarios described below.

### 2.2. Simulation scenarios

Different simulation scenarios are defined to assess how SMCs influence vessel-scale pressure-diameter responses and the resulting organ-scale pressure-flow curve. Four scenarios are considered to isolate the effects of SMC activation, structural heterogeneity in SMC abundance, and functional heterogeneity in SMC pressure-calcium behavior:

#### Passive reference scenario

In this scenario, intracellular calcium is fixed at its baseline value throughout the simulation, eliminating pressure-dependent SMC activation. This approach preserves the baseline mechanical state of the vessel wall while eliminating active myogenic feedback, providing a consistent reference against which active SMC responses can be evaluated.

#### Homogeneous SMC scenario

Here, SMCs are assumed homogeneous in both abundance and behavior throughout the vascular network. Consequently, all vessels are assigned the same SMC volume fraction and pressure-calcium relationship. To describe the latter, any of the four experimental pressure-calcium relationships reported by Cipolla et al.^8^ and Nystoriak et al.^10^ for cerebral arteries and arterioles (Figure 2A) could be used. In the present study, the cerebral arteriole data of Nystoriak et al. ^10^ were selected and fitted by

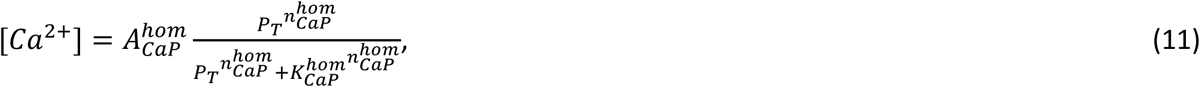

with 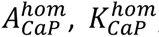, and 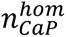 determined through a nonlinear least-squares fitting and *P_T_* the transmural pressure. Results obtained using the three alternative pressure-calcium relationships (Figure 2A) are provided in Supplementary Material S3.

**Figure 2.**
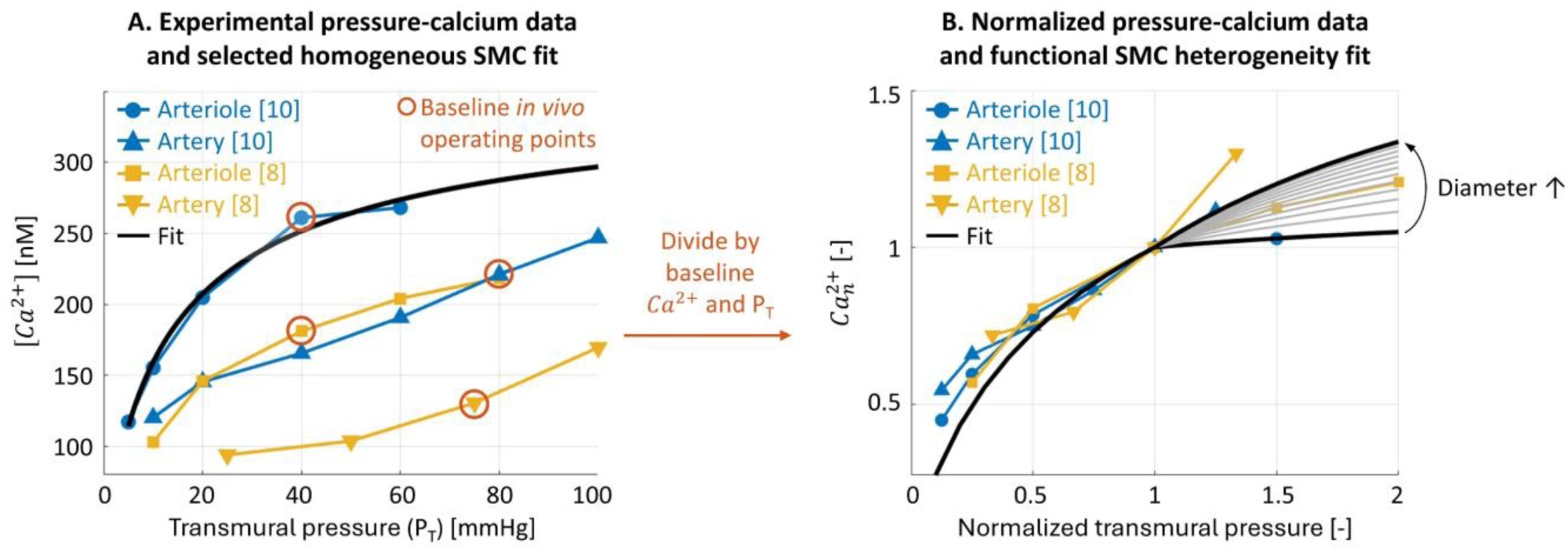
Pressure-calcium relationships used to define the homogeneous and functional SMC heterogeneity scenarios. (A) Experimental pressure-calcium relationships from Nystoriak et al.^10^ (parenchymal arterioles: blue circles, middle cerebral arteries: blue upward triangles) and Cipolla et al.^8^ (parenchymal arterioles: yellow squares, middle cerebral arteries: yellow downward triangles). The black curve denotes the fit to the Nystoriak et al.^10^ parenchymal arteriole dataset selected to describe the homogeneous SMC scenario (Equation 11). Red circles indicate the baseline in vivo operating points of each dataset: Nystoriak et al.^10^ arterioles (40mmHg, 261nM) and arteries (80mmHg, 221nM), Cipolla et al.^8^ arterioles (40mmHg, 181nM) and arteries (75mmHg, 131nM). (B) Pressure-calcium relationships after normalizing the data of panel A by their baseline in vivo pressure and calcium values (red circles). The black and grey curves represent the diameter-dependent normalized pressure-calcium formulation used in the functional SMC heterogeneity scenario (Equation 12). The diameter dependence was described based on the data from Nystoriak et al.^10^, as it explicitly reports the diameters of the tested vessels.

#### Structural SMC heterogeneity scenario

In this scenario, SMC pressure-calcium behavior is assumed homogeneous, while heterogeneity in SMC abundance is introduced through variations in SMC volume fraction. Experimental evidence suggests reduced SMC coverage in smaller, more distal vessels^5^, but quantitative data describing its variation across the vascular tree are lacking. Consequently, vessels are just partitioned into two diameter-based groups, with the smaller-vessel group assigned half the SMC volume fraction of the larger-vessel group. In both groups, the remaining wall volume is distributed between elastin and collagen under a fixed fraction. The diameter threshold separating both groups is set at 70 µm, as explained in Section 2.4.

#### Functional SMC heterogeneity scenario

SMC abundance is assumed homogeneous throughout the vascular network, while heterogeneity in SMC behavior is introduced through vessel-size-dependent pressure-calcium relationships. Because experimental pressure-calcium data are only available for two vessel sizes^8,10^, directly defining separate relationships for all vessel sizes in the network is not feasible. To obtain a unified formulation while retaining vessel-size dependence, the experimental datasets are normalized by their respective baseline *in vivo* transmural pressure and intracellular calcium values (circles in Figure 2A). In this normalized representation, the datasets exhibit improved agreement, enabling their description by a common Hill-type formulation in which selected parameters vary continuously with vessel diameter (Figure 2B):

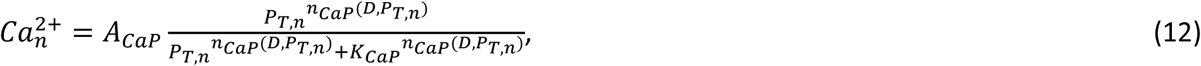

where *P_T_*_,*n*_ denotes normalized transmural pressure, *A_CaP_* is a scaling constant, *K_CaP_* the half-activation constant, and *n_CaP_*(*D*, *P_T_*_,*n*_) a diameter-and pressure-dependent Hill exponent.

The corresponding intracellular calcium concentration is obtained as

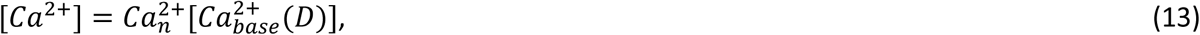

with 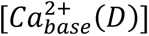 the diameter-dependent baseline calcium concentration. The equations describing *n_CaP_*(*D*, *P_T_*_,*n*_) and 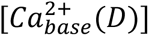 are parameterized analogous to Equation 2 (Supplementary Material S1.3.), using the Nystoriak et al. dataset^10^, as this dataset explicitly reports the diameters of the tested vessels.

### 2.3. Parameter calibration and sensitivity analysis

Calibration and sensitivity analysis are performed to define the physiological operating point of the model and assess the robustness of model predictions to parameter uncertainty, respectively. Given the high dimensionality of the parameter space, sensitivity analysis is conducted using the Morris method (with p=4 and 40 trajectories) which enables efficient global screening^35–37^. In this approach, elementary effects are computed for each parameter by evaluating the change in model output resulting from perturbations in parameter values. The output of interest is the relative cerebral blood flow at each level of cerebral perfusion pressure applied to the organ scale vascular network. Parameter influence is quantified by the mean of the absolute elementary effects (µ*) and the standard deviation of the elementary effects (σ), reflecting overall sensitivity and nonlinear or parameter interaction effects, respectively^35^.

Calibrated parameter values, sensitivity ranges, and used sources are summarized in Table 1. Parameters specific to the homogeneous SMC scenario are not assigned ranges because sensitivity analysis is performed only for the most complex scenario (functional SMC heterogeneity). Where possible, parameters are calibrated using human data. The source columns in Table 1 indicate both the source(s) and a numerical code specifying how the sources were used to set the parameter value or range: (1) The exact value from the source is used, (2) A representative value in the range of the observations of the source(s) is used, (3) The parameter is fitted to experimental data of the source, (4) The parameter range is inferred from the variability across different sources, (5) An explanation is provided in Supplementary Material S2. When no quantitative information on parameter variability is available, sensitivity ranges are set to ±25 % of the calibrated value^37^.

**Table 1.** Overview of parameter values for calibration and parameter ranges for the sensitivity analysis. Next to the sources a code is used to indicate how the value or range was obtained from the source: (1) The exact value from the source is used, (2) A representative value in the range of the observations of the source(s) is used, (3) The parameter is fitted to experimental data of the source, (4) The parameter range is inferred from the variability across different sources, (5) An explanation is provided in Supplementary Material S2. ‘-‘ indicates no source was available. In these cases the range is set as ±25 % relative to the calibrated baseline value. ‘NA’ indicates ‘Not applicable’.

| Parameter | Value | Source | Range | Source |
| --- | --- | --- | --- | --- |
| ORGAN SCALE |  |  |  |  |
| $D_{in}$ | 2860 $\mu\text{m}$ | [<sup>41</sup>] (1) | 2520 to 3200 $\mu\text{m}$ | [<sup>41</sup>] (1) |
| $D_c$ | 8 $\mu\text{m}$ | [<sup>22,24,42</sup>] (1) | 5 to 10 $\mu\text{m}$ | [<sup>12,22,42,43</sup>] (4) |
| $\lambda_{a/c}$ | 1.25 | [<sup>12,23</sup>] (2) | 1 to 2 | [<sup>12,23</sup>] (2) |
| $\lambda_{v/a}$ | 2.1 | [<sup>22</sup>] (2) | 1.2 to 2.15 | [<sup>22,25,42</sup>] (2) |
| $D_{M,l}$ | 500 $\mu\text{m}$ | [<sup>20</sup>] (1) | $\pm 25\%$ | - |
| $n_{M,s}$ | 3.98 | [<sup>21</sup>] (2) | 3.78 to 4.18 | [<sup>20,21</sup>] (2) |
| $n_{M,l}$ | 2.5 | [<sup>20</sup>] (1) | 2.3 to 2.7 | [<sup>20</sup>] (2) |
| $S_M$ | -0.3 | [<sup>20</sup>] (3) | -0.7 to 0.1 | [<sup>20</sup>] (2,3) |
| $N_c$ | 10 | [<sup>22,44-46</sup>] (5) | $\pm 25\%$ | - |
| $\lambda_{L/D,a}$ | 20 | [<sup>22,42</sup>] (2) | 10 to 35 | [<sup>22,42</sup>] (2) |
| $\lambda_{L/D,c}$ | 125 | [<sup>22</sup>] (1) | $\pm 25\%$ | - |
| $\lambda_{L/D,v}$ | 13 | [<sup>22</sup>] (2) | 5 to 20 | [<sup>22</sup>] (2) |
| $\lambda_{t/D,a}$ | 0.15 | [<sup>22,25</sup>] (2) | 0.1 to 0.2 | [<sup>22,25</sup>] (2) |
| $\lambda_{t/D,c}$ | 0.09 | [<sup>22,25</sup>] (2) | 0.0625 to 0.125 | [<sup>22,25</sup>] (2) |
| $\lambda_{t/D,v}$ | 0.05 | [<sup>22,25</sup>] (2) | 0.03 to 0.1 | [<sup>22,25</sup>] (2) |
| $\mu_{plasma}$ | 1.125 cP | [<sup>47</sup>] (1) | 1 to 1.6 | [<sup>7,47,48</sup>] (4) |
| $H_D$ | 0.42 | [<sup>7</sup>] (1) | 0.4 to 0.45 | [<sup>7,47,48</sup>] (4) |
| $P_{in}$ | 85 mmHg | [<sup>12,17,22,49</sup>] (2) | 75 to 95 mmHg | [<sup>12,17,22,49</sup>] (4) |
| $MAP - P_{in}$ | 5 mmHg | [<sup>22,50</sup>] (2) | 3 to 7 mmHg | [<sup>22,49,50</sup>] (2) |
| $P_{ICP}$ | 10 mmHg | [<sup>11,24</sup>] (1) | 7 to 15 mmHg | [<sup>51</sup>] (1) |
| $P_{out} - P_{ICP}$ | 4 mmHg | [<sup>17,22</sup>] (2) | 0 to 6 mmHg | [<sup>22</sup>] (2) |
| VESSEL SCALE |  |  |  |  |
| $\phi_{el}$ | 0.05 | [<sup>52</sup>] (2) | Reparametrized using elastin-collagen fraction, see below. (5) | |
| $\phi_{col}$ | 0.15 | [<sup>52</sup>] (2) | | |
| $\theta_{e/c} = \frac{\phi_{el}}{\phi_{el} + \phi_{col}}$ | 0.25 | $\frac{0.05}{0.05 + 0.15}$ | 0.17 to 0.33 | [<sup>52</sup>] (2,5) |
| $\phi_{SMC}$ | 0.8 | [<sup>52</sup>] (2) | 0.7 to 0.9 | [<sup>52</sup>] (2) |
| MICROSTRUCTURAL SCALE |  |  |  |  |
| $g_{zz,el}$ | 1.17 | [<sup>53</sup>] (1) | 1.1 to 1.3 | [<sup>54,55</sup>] (4) |
| $g_{col}$ | 1.1 | [<sup>29</sup>] (1) | 1.05 to 1.15 | [<sup>56,57</sup>] (4) |
| $C_{10,el}$ | 0.08 MPa | [<sup>58</sup>] (3) | 0.05 to 0.16 MPa | [<sup>59</sup>] (1) |
| $k_{1,col}$ | 0.73 MPa | [<sup>58</sup>] (3) | $\pm 25\%$ | - |
| $k_{2,col}$ | 0.91 | [<sup>58</sup>] (3) | $\pm 25\%$ | - |
| $\kappa_{col}$ | 0.26 | [<sup>58</sup>] (3) | 0 to 1/3 | [<sup>60</sup>] (1) |
| $\alpha_{col}$ | 1.7° | [<sup>58</sup>] (3) | 0 to 25° | [<sup>29,60,61</sup>] (4) |
| $\mu_{SMC}$ | 0.31 MPa | [<sup>62</sup>] (2) | $\pm 25\%$ | - |
| $\kappa_{SMC}$ | 1.16 MPa | [<sup>62</sup>] (2) | $\pm 25\%$ | - |
| $\alpha_{SMC}$ | 0° | [<sup>29</sup>] (1) | 0 to 20° | [<sup>29,63</sup>] (4) |
| $C_{10,c}$ | 62 kPa | [<sup>64</sup>] (1) | ±25% | - |
| $C_{10,v}$ | 65 kPa | [<sup>64</sup>] (1) | ±25% | - |
| MOLECULAR SCALE: HAI-MURPHY |  |  |  |  |
| $k_2=k_5$ | $0.5 \text{ s}^{-1}$ | [<sup>29-31</sup>] (1) | ±25% | - |
| $k_3$ | $0.4 \text{ s}^{-1}$ | [<sup>29-31</sup>] (1) | ±25% | - |
| $k_4$ | $0.1 \text{ s}^{-1}$ | [<sup>29-31</sup>] (1) | ±25% | - |
| $k_7$ | $0.01 \text{ s}^{-1}$ | [<sup>29-31</sup>] (1) | ±25% | - |
| $k_{1,max}$ | $0.5 \text{ s}^{-1}$ | (5) | ±25% | - |
| $K_{k1}$ | 178 nM | [<sup>29,30</sup>] (1) | ±25% | - |
| $n_{k1}$ | 2 | [<sup>29,30,65</sup>] (1) | ±25% | - |
| $CaM_{max}$ | Optimized within literature range | [<sup>33</sup>] (5) | NA (5) | NA (5) |
| $K_{CaM}$ | Optimized within literature range | [<sup>33</sup>] (5) | NA (5) | NA (5) |
| $k_{1,base}$ | $0.2 \text{ s}^{-1}$ | [<sup>30</sup>] (2,5) | ±25% | - |
| $n_{CaM}$ | 2.5 | [<sup>32,33</sup>] (2) | 1 to 4 | [<sup>32,33</sup>] (2) |
| MOLECULAR SCALE: PRESSURE-CALCIUM |  |  |  |  |
| $A_{CaP}^{hom}$ | 359.01 nM | [<sup>10</sup>] (3) | NA | NA |
| $K_{CaP}^{hom}$ | 13.37 mmHg | [<sup>10</sup>] (3) | NA | NA |
| $n_{CaP}^{hom}$ | 0.78 | [<sup>10</sup>] (3) | NA | NA |
| $A_{CaP}$ | Defined by $K_{CaP}$ and $n_{CaP}$ under fitting constraint. (5) | | | |
| $K_{CaP}$ | 1 | [<sup>10</sup>] (3) | ±25% | - |
| $n_{CaP,L}$ | 0.8 | [<sup>10</sup>] (3) | ±25% | - |
| $D_{CaP,s}$ | 63 $\mu\text{m}$ | [<sup>10,40</sup>] (5) | 53 to 73 $\mu\text{m}$ | [<sup>10,40</sup>] (5) |
| $D_{CaP,l}$ | 2860 $\mu\text{m}$ | [<sup>10,41</sup>] (5) | 2520 to 3200 $\mu\text{m}$ | [<sup>10,41</sup>] (5) |
| $n_{CaP,H,s}$ | 0.14 | [<sup>10</sup>] (3) | ±25% | - |
| $n_{CaP,H,l}$ | 1.01 | [<sup>10</sup>] (3) | ±25% | - |
| $S_N$ | 0.5 | (5) | 0.5 to 1.5 | (5) |
| $Ca_{base,s}$ | 261 nM | [<sup>10</sup>] (1) | 241 to 281 nM | [<sup>10</sup>] (2) |
| $Ca_{base,l}$ | 221 nM | [<sup>10</sup>] (1) | 201 to 241 nM | [<sup>10</sup>] (2) |
| $S_{Ca}$ | -0.05 | (5) | -0.05 to 2.05 | (5) |

### 2.4. Comparison with experimental data

Due to the limited availability of *in vivo* cerebral autoregulation data in humans across wide pressure ranges, model predictions are compared against experimental piglet data^5^. In these experiments, cerebral perfusion pressure was varied quasi-statically by mechanical manipulation while cerebral blood flow and vessel diameters were monitored through a cranial window. Averaging responses across observed vessels and animals (10 hypotension and 10 hypertension) yielded aggregate pressure-flow and pressure-diameter relationships. In addition, pressure-diameter curves were constructed for three vessel size classes: diameters below 40 µm, between 40 and 70 µm, and between 70 and 123 µm, to see whether vessel-size-specific responses could be observed. Consistent with this experimental classification, a diameter threshold of 70 µm is used in the structural SMC heterogeneity scenario (see Section 2.2), such that reduced SMC abundance (*φ_el_*=0.15, *φ_col_*=0.45, *φ_SMC_*=0.4) is assigned only to vessels in the two smaller diameter classes.

As the model was primarily calibrated to human values, comparison with piglet experiments requires accounting for interspecies differences. While the cerebral vasculature of piglets is similar to that of humans^38^, clear differences exist in vessel dimensions and baseline pressures. A limited set of parameters is therefore adapted using reported values or scaling relations, as shown in Table 2.

**Table 2.** Overview of parameter adaptations applied to enable comparison between the human-calibrated model and experimental piglet data.

| Parameter | Value for humans | Value for piglets | Source |
| --- | --- | --- | --- |
| $D_{in}$ | 2860 $\mu\text{m}$ | 775 $\mu\text{m}$ | [<sup>66-68</sup>] |
| $D_c$ | 8 $\mu\text{m}$ | 6.3 $\mu\text{m}$ | [<sup>22,40,66</sup>] |
| $D_{M,l}$ | 500 $\mu\text{m}$ | 200 $\mu\text{m}$ | [<sup>20,40,66-68</sup>] |
| $P_{in}$ | 85 mmHg | 70 mmHg | [<sup>5,69</sup>] |
| $P_{ICP}$ | 10 mmHg | 5 mmHg | [<sup>5,69</sup>] |
| $P_{out} - P_{ICP}$ | 4 mmHg | 2.5 mmHg | [<sup>5,69</sup>] |
| $D_{CaP,s}$ | 63 $\mu\text{m}$ | 50 $\mu\text{m}$ | [<sup>10,40,66</sup>] |

## 3. Results

### 3.1. Vessel-size-dependent pressure-diameter responses

The effect of SMC activation and heterogeneity on vessel-scale behavior is examined by comparing pressure-diameter responses across vessel size classes for the different simulation scenarios (Figure 3). To assess which SMC representation best reproduces physiological vessel behavior, the simulated responses are evaluated against experimentally observed size-dependent trends, which show that large vessels maintain constriction until higher pressures than smaller vessels^5^, exhibit the greatest dilation during initial pressure reductions^9^, and attain a lower maximal dilation under further pressure reductions^5^.

**Figure 3.**
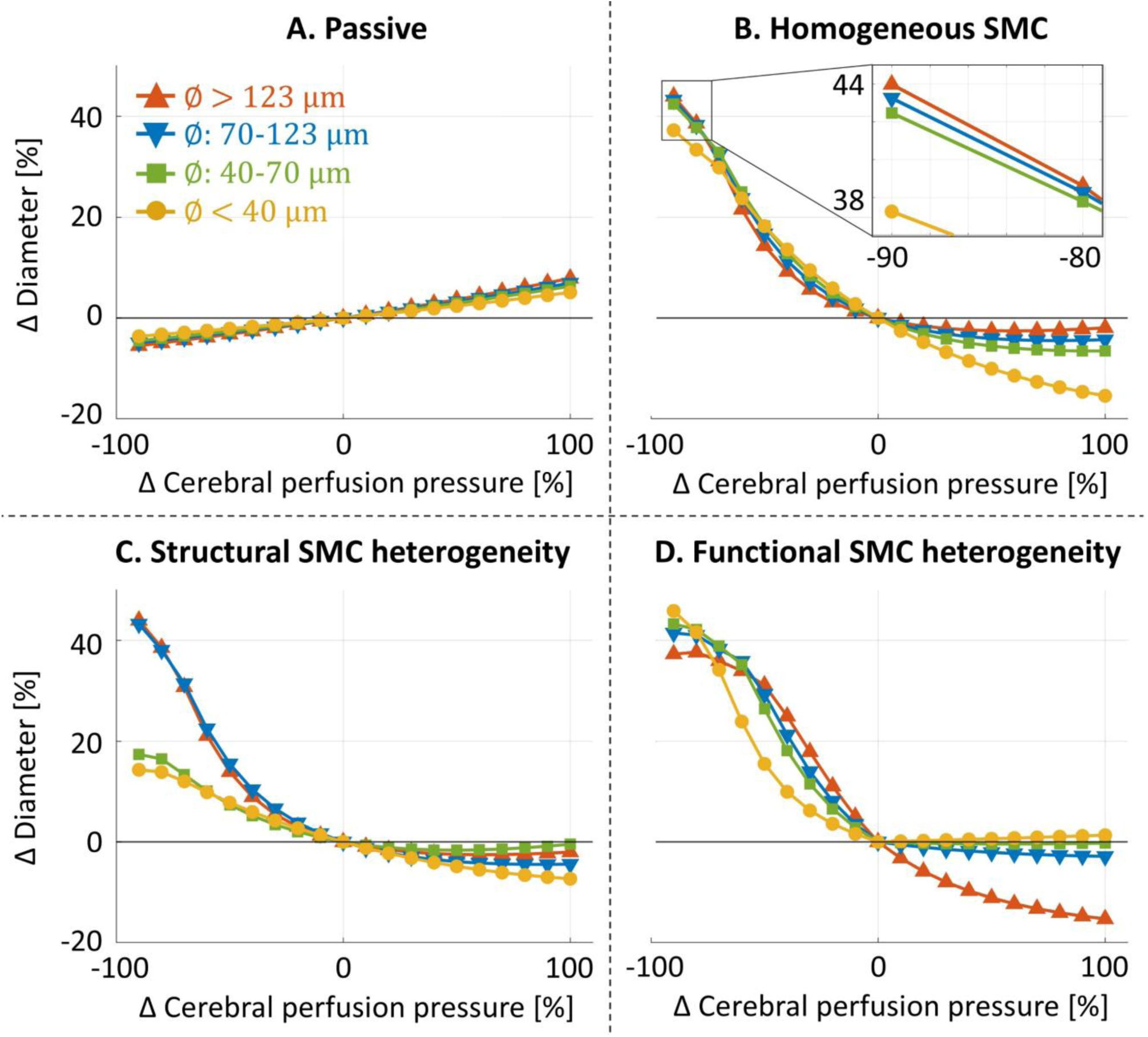
Vessel diameter (relative to baseline) as a function of cerebral perfusion pressure (relative to baseline) for the four simulation scenarios, with responses grouped by vessel size class. Panels A to D correspond to the passive reference, homogeneous SMC, structural SMC heterogeneity, and functional SMC heterogeneity scenarios, respectively. Within each panel, the different curves represent the average pressure-diameter response of all vessels within a given size class: diameters <40 µm in yellow circles, 40-70 µm in green squares, 70-123 µm in blue downward triangles, >123 µm in red upward triangles.

In the passive reference scenario (Figure 3A), vessel diameters decrease at reduced pressures and increase at elevated pressures, indicating purely passive behavior. Differences between vessel size classes are minimal, with only a slight relative reduction in deformation in the smaller vessels.

Introducing homogeneous SMC activation (Figure 3B) restores autoregulatory behavior, with vessels dilating at reduced pressures and constricting at elevated pressures. However, the predicted size dependence is opposite to that observed experimentally. During the initial decrease in pressure, the smallest vessels exhibit the largest relative dilation and as pressure decreases further, large vessels progressively dilate more and eventually show the greatest overall dilation. At elevated pressures, all vessels initially constrict but as pressure increases further, larger vessels start to increase in diameter again. Vessels in the >123 µm group reach their maximal constriction (-2.56%) at +60% CPP, followed by the 70-123 µm and 40–70 µm groups at +80% (-4.48%) and +90% ΔCPP (-6.54%), respectively, while the smallest vessels continue to decrease in diameter over the full pressure range considered.

In the structural SMC heterogeneity scenario (Figure 3C) reducing the SMC volume fraction in vessels below 70 µm, causes them to dilate less at reduced pressures and constrict less at elevated pressures, compared to the homogenous scenario (Figure 3B). The 40-70 µm group also reaches maximal constriction earlier, attaining a minimum relative diameter of -1.65% at +50% ΔCPP. While this brings the high-pressure response closer to the experimentally observed size-dependent trends, it increases discrepancies at reduced pressures^5,9^.

In the functional SMC heterogeneity scenario (Figure 3D), the smallest vessels dilate the least during the initial pressure decrease, show the greatest maximal dilation under further pressure reductions, and are the first to lose active constriction as pressure increases, all consistent with experiments^5,9^.

### 3.2. Organ-scale pressure-diameter and pressure-flow responses

The effect of SMC activation and heterogeneity on organ-scale autoregulation is examined by comparing aggregate pressure-flow and pressure-diameter curves for the different simulation scenarios among each other and to experiments^5^ (Figure 4). Experimentally, vessels dilate at reduced pressures and constrict at elevated pressures before transitioning to passive dilation. The response is asymmetric, with greater diameter changes during hypotension than hypertension, resulting in near-constant flow at initial pressure reductions but an immediate increase in flow at elevated pressures^5^. Note that because the experimental measurements are limited to vessels visible through the cranial window, corresponding model curves are obtained by averaging over the same vessel sizes rather than over the full network to allow for fair comparison.

**Figure 4.**
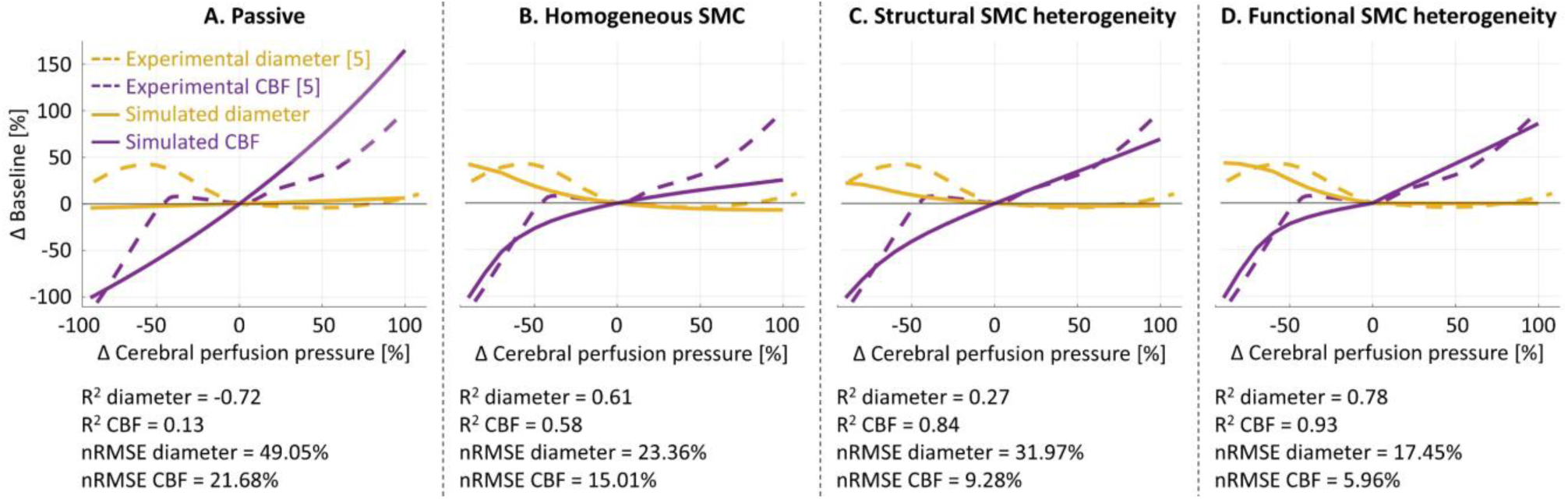
Organ-scale pressure-flow and pressure-diameter curves (relative to baseline) for the four simulation scenarios, comparing model predictions (full lines) to experimental data ^5^ (dashed lines), with cerebral blood flow (CBF) shown in purple and vessel diameter in yellow. Panels A to D correspond to the passive reference, homogeneous SMC, structural SMC heterogeneity, and functional SMC heterogeneity scenarios, respectively. Quantitative agreement between model and experiments is reported using the coefficient of determination (R^2^) and the normalized root mean square error (nRMSE).

In the passive reference scenario (Figure 4A), vessel diameters decrease at reduced pressures and increase at elevated pressures, resulting in no stabilization of cerebral blood flow. This mismatch between model predictions and experimental observations^5^ is reflected in poor agreement metrics, with a negative coefficient of determination for diameter (R^2^ = -0.72), a low coefficient for CBF (R^2^ = 0.13), and large normalized root mean square errors for both (nRMSE = 49.05% for diameter and 21.68% for flow).

Introducing homogeneous SMC activation (Figure 4B) results in vessels dilating at reduced pressures and constricting at elevated pressures, leading to partial stabilization of CBF around baseline. However, substantial discrepancies with experimental data remain, including a more symmetric pressure-diameter and pressure-flow response than observed experimentally. Consequently, agreement metrics remain moderate with R^2^ diameter = 0.61, R^2^ CBF = 0.58, nRMSE diameter = 23.36% and nRMSE CBF = 15.01%.

In the structural SMC heterogeneity scenario (Figure 4C), vessels exhibit on average less dilation at reduced pressures and less constriction at elevated pressures than in the homogeneous scenario (Figure 4B), reaching maximal constriction within the tested pressure range (at +60% ΔCPP), before passively dilating. As a result, CBF shows larger deviations from baseline. While the altered response improves agreement with experimental observations at elevated pressures, it worsens agreement at reduced pressures. Overall, the net effect is improved CBF predictions (R^2^ = 0.84, nRMSE = 9.28%) but worse diameter predictions (R^2^ = 0.27, nRMSE = 31.97%).

In the functional SMC heterogeneity scenario (Figure 4D), the pressure-diameter response becomes more asymmetric, with stronger dilation at reduced pressures than constriction at elevated pressures. This asymmetry is mirrored in the pressure-flow relationship, where flow increases more sharply at elevated pressures than it decreases at reduced pressures. As a result, this scenario shows the best overall agreement with experimental data^5^, yielding the highest coefficients of determination (R^2^ diameter = 0.78, R^2^ CBF = 0.93) and lowest errors (nRMSE diameter = 17.45%, nRMSE CBF = 5.96%) among all scenarios.

### 3.3. Sensitivity analysis

Parameter sensitivity is assessed for the most elaborate scenario (functional SMC heterogeneity) by quantifying the influence of each parameter on relative CBF across the pressure range (purple full line shown in Figure 4D). Figure 5 summarizes the parameter sensitivities using scatter plots of µ* and σ averaged over the full (A), reduced (B), and elevated (C) pressure ranges. Parameters located in the upper right region of these plots simultaneously combine a strong influence on model output (µ*) with pronounced nonlinearity or interaction with other parameters (σ)^35,37^, indicating a dominant role in governing system behavior. For each scatter plot (A-C) the ten most influential parameters are highlighted. Their ranking is based on the geometric mean of the averaged µ* and σ values after normalizing them by the largest average µ* and σ across all parameters.

**Figure 5.**
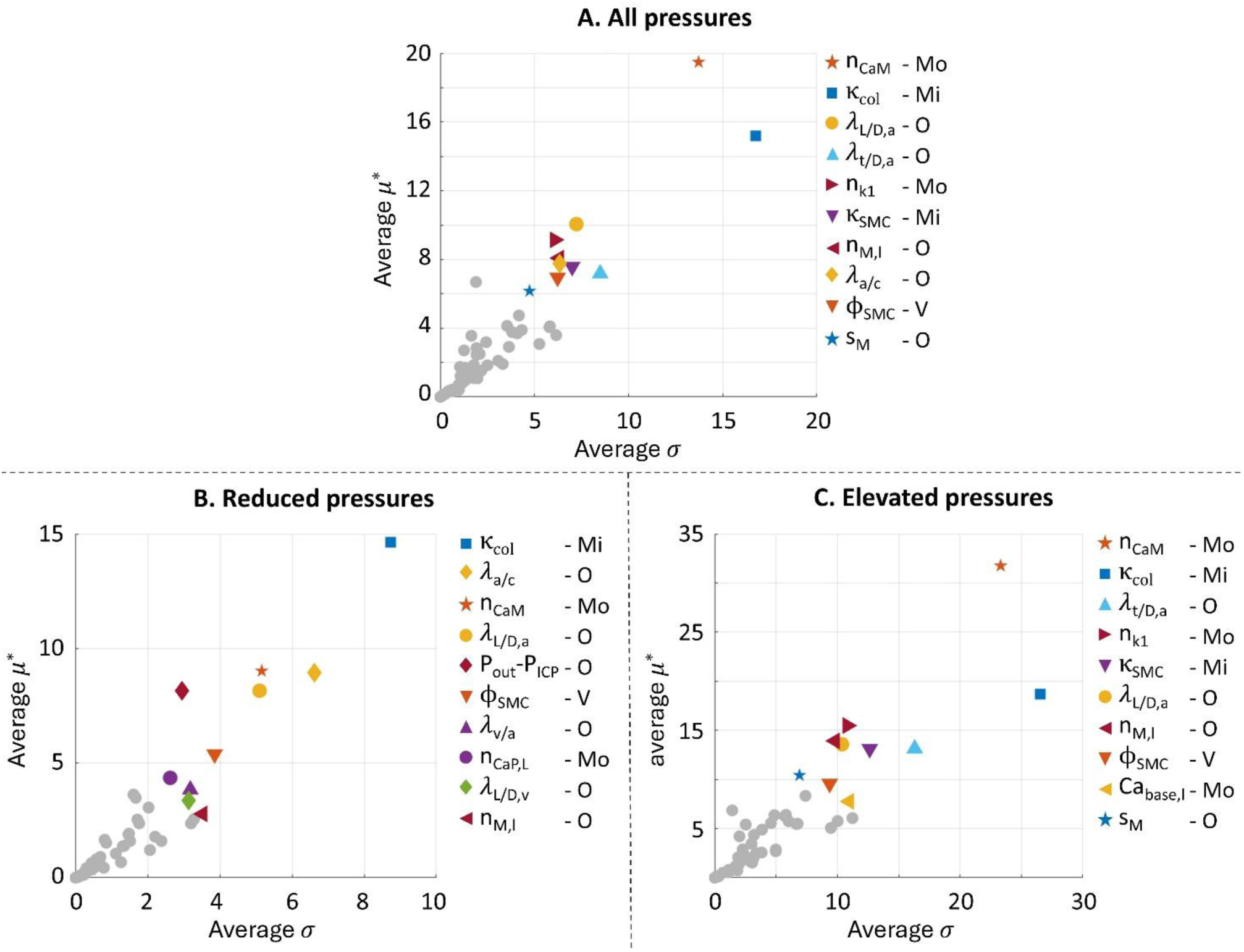
Sensitivity analysis results for the functional SMC heterogeneity scenario based on the Morris method, showing parameter influence on relative cerebral blood flow. For each parameter, µ* (parameter influence) and σ (parameter nonlinearity and/or interaction with other parameters) are first computed at individual pressure levels along the pressure-flow curve and subsequently averaged over (A) the full, (B) reduced, and (C) elevated pressure ranges. Scatter plots display the averaged μ* versus σ values for all parameters. The ten parameters with the highest combined effect (geometric mean of normalized μ* and σ) are highlighted. Remaining parameters are shown as grey circles. Next to each parameter it is indicated whether the parameter relates to the organ (O), vessel (V), microstructural (Mi) or molecular (Mo) scale.

Across the full pressure range (Figure 5A), the top ten parameters span all model scales (organ: O, vessel: V, microstructure: Mi, molecular: Mo). This pattern is also observed in the reduced (Figure 5B) and elevated pressure (Figure 5C) regimes, but the balance across scales differs. For example, organ scale parameters are more strongly represented at reduced pressures, accounting for six of the top ten parameters (including three of the top five), compared with four of ten (and one of five) at elevated pressures. A more detailed view of the pressure-dependent evolution of parameter importance is provided in Supplementary Material S4.

## 4. Discussion

We developed a multi-scale model of cerebral autoregulation to investigate how SMC activation and heterogeneity shape cerebral autoregulation across scales. In particular, we examined how these mechanisms shape vessel-scale pressure-diameter responses and how their size-dependent behavior translates into the global pressure-flow relationship at the organ scale.

### 4.1. Vessel-size-dependent pressure-diameter responses

The results indicate that SMC activity is necessary but not sufficient to explain the experimentally observed size-dependent pressure-diameter responses. The necessity of SMC activity is evident when comparing the passive scenario (Figure 3A) to the homogeneous SMC activation scenario (Figure 3B), as the latter transforms the model from a purely elastic mechanical system into one capable of myogenic diameter regulation. This finding is consistent with the established role of SMCs in cerebral autoregulation^1,7,34^. However, homogeneous SMC activation fails to reproduce the observed vessel-size-dependent behavior, demonstrating that SMC activity alone is insufficient. This conclusion is independent of the selected experimental pressure-calcium relationship to describe the homogeneous response (Section 2.2) as shown in Supplementary Material S3.

Structural SMC heterogeneity can contribute to vessel-size-dependent behavior, but it cannot explain the observed responses across the full pressure range. By altering the amount of contractile tissue available for force generation, differences in SMC abundance scale the magnitude of the active response. Reducing SMC abundance in smaller vessels (Figure 3C) shifts their behavior toward a more passive response, thereby changing their relative response magnitude compared to the other vessel size classes. However, because the introduced heterogeneity is a fixed scaling, independent of pressure, it cannot reproduce the experimentally observed reversal in the size-dependent behavior, whereby small vessels exhibit the greatest maximal dilation during pressure reductions while large vessels maintain constriction over a broader pressure range during pressure elevations^5^. This limitation would persist even if data became available to replace the current two-group calibration (Section 2.2) with a continuous variation in SMC abundance along the vascular tree, as such a refinement would only modify the degree of scaling, not its pressure independence.

In contrast, functional SMC heterogeneity introduces vessel-size-dependent differences that are inherently pressure dependent. This arises because the vessel-size-dependent pressure-calcium responses underlying this heterogeneity differ not only in magnitude but also in how their local slope, and thus their pressure sensitivity, varies across the pressure range (Figure 2A). Consequently, the vessel size class exhibiting the greatest pressure sensitivity shifts from small vessels below baseline pressure to large vessels above baseline pressure. In the normalized domain used to model the pressure-calcium responses (Equation 12, Figure 2B), this behavior is encoded by the combination of normalized slopes and the vessel-size-specific ratio of baseline intracellular calcium to baseline transmural pressure, which is larger in smaller vessels.

Beyond introducing pressure-dependent differences in pressure sensitivity, functional SMC heterogeneity influences how calcium signals are translated into SMC activation. The calcium-to-activation mechanism itself, represented by the Ca^2+^-*k*_1_ relationship, is assumed to be identical across vessel sizes (Equations 9 and 10), consistent with the observation by Cipolla et al.^8^ that the contractile apparatus does not exhibit size-dependent differences. Size dependency emerges because vessels occupy distinct operating points along this common nonlinear Ca^2+^-*k*_1_ curve. These operating points are determined by the vessel-size-dependent baseline intracellular calcium concentrations set by the vessel-size-dependent pressure-calcium relationships of the functional SMC heterogeneity scenario^8,10^. In the functional heterogeneity scenario, smaller vessels operate at higher baseline calcium levels, positioning them closer to the upper saturation limit. This increases their dilation reserve due to their greater distance to the lower saturation limit, while simultaneously reducing their Ca^2+^-to-*k*_1_ gain during further calcium increases as the local Ca^2+^-*k*_1_ slope declines near saturation. In contrast, the common pressure-calcium relationship in the homogeneous scenario, causes larger vessels, which experience higher baseline pressures, to operate at higher baseline calcium levels, producing the opposite size-dependent pattern.

Together, the differences introduced by functional SMC heterogeneity allow the model to reproduce the experimentally observed vessel-size-dependent pressure-diameter responses^5,9^ (Figure 3D). The reversal in size-dependent trends reported by Klein et al.^5,9^ follows from the combined pressure-dependent effects on pressure sensitivity and calcium-to-*k*_1_ coupling described above. Below baseline pressure, smaller vessels exhibit greater pressure sensitivity, a higher Ca^2+^-to-*k*_1_ gain, and greater dilation reserve, resulting in the greatest maximal dilation. Above baseline pressure, larger vessels exhibit greater pressure-to-*k*_1_ sensitivity and greater constriction reserve, allowing them to maintain active constriction over a broader pressure range. Despite the greater dilation capacity of smaller vessels, large vessels exhibit a greater dilation response during the first pressure reductions from baseline, consistent with the findings of Kontos et al.^9^. This likely reflects that the larger absolute pressure reduction experienced by large vessels outweighs their lower pressure-to-*k*_1_ sensitivity at this point in the pressure range.

Although functional SMC heterogeneity captures the experimentally observed size-dependent trends, the magnitude of differences between vessel groups remains smaller than observed experimentally, particularly at reduced pressures. In the piglet experiments, maximal dilation differs by approximately 20% between consecutive vessel groups (<40 µm, 40–70 µm, and 70-123 µm)^5^, while the model predicts differences of only around 2% between these groups. A potential explanation is that the model only includes the myogenic response. Although it is commonly described as the dominant mechanism in CBF regulation^1,39^, the other mechanisms active *in vivo*, such as the endothelial response, may influence size-dependent differences as well.

### 4.2. Organ-scale pressure-diameter and pressure-flow responses

The organ scale results (Figure 4) support the hypothesis of Klein et al.^5^ that vessel-size-dependent pressure-diameter responses contribute to the characteristic pressure-flow relationship. Across the four simulation scenarios, improved agreement with the experimentally observed vessel-size-dependent pressure-diameter responses (Section 3.1) is consistently accompanied by improved agreement with the aggregate pressure-diameter and pressure-flow responses (Section 3.2). This correspondence is particularly evident for the structural SMC heterogeneity scenario, where the improved agreement with vessel-size-dependent responses at elevated pressures, but deteriorated agreement at reduced pressures, is mirrored by corresponding improvements and deteriorations in the aggregate autoregulatory curves. The remaining discrepancies between the functional SMC heterogeneity and experimental observations^5^ (Figure 4D) may reflect the model’s restriction to the myogenic mechanism, as other mechanisms likely contribute *in vivo*, as also discussed for the vessel-scale results in Section 4.1.

### 4.3. Sensitivity analysis

The sensitivity analysis presented in Figure 5 shows that the most influential parameters span all model scales, highlighting that model behavior depends on contributions from all levels of the system rather than being dominated by a single scale. The balance between scales, however, varies with pressure, with organ-scale parameters relatively more influential at reduced pressures. A potential explanation is that, as pressure decreases, vessels approach their maximal dilation (Figure 3D), limiting active SMC control and causing the response to become increasingly governed by network structure and pressure distribution. This interpretation is supported by the results in Supplementary Material S4, which show that organ scale parameters gain importance as pressure decreases further, while the influence of molecular parameters diminishes. At elevated pressures, large vessels have not yet reached their maximal constriction (Figure 3D), meaning that active SMC control still plays an important role. As a result, there is no transition toward organ scale dominance with pressure and the overall influence of organ scale parameters remains lower compared to the reduced pressure regime.

### 4.4. Limitations

As with any computational model, physiological realism was balanced against model interpretability and computational efficiency. Because the aim of this study was to investigate how SMC heterogeneity shapes autoregulation during slow changes in perfusion pressure, the model focused on the steady-state myogenic response, neglecting possible contributions from other CBF regulation pathways^1^. Furthermore, to facilitate comparison between vessel size classes, the organ-scale vasculature was represented as a symmetric bifurcating tree without anastomotic connections such that each vessel size class corresponds to a unique branch level and loading condition rather than to a collection of same sized vessels distributed across different network positions. While beneficial for defining and comparing vessel size classes, this simplification reduces anatomical realism and limits the applicability of the model to patient-specific simulations. Future work could address this limitation by incorporating patient-specific vascular networks and embedding the cerebral circulation within a whole-body model, thereby also removing the assumptions of constant intracranial and outlet pressures.

A further limitation stems from incomplete experimental knowledge and quantitative data regarding several aspects of cerebral autoregulation. Capillaries were represented as passive elements, neglecting potential active contributions from pericytes^1,7^, as their role in cerebral autoregulation remains insufficiently quantified to include reliably^1^. The vessel-size-dependent pressure-calcium responses were extrapolated beyond the experimentally measured pressure range^8,10^ to describe behavior across the full pressure range considered in this study. Furthermore, to enable a complete description of all model scales, measurements from multiple species were incorporated where necessary. Although species-specific differences were accounted for where possible, for example by scaling the vessel diameter *D_CaP_*_,*s*_ from rat^10^ to human^40^ (Table 1 and Supplementary Material S2), uncertainty associated with interspecies differences in vascular regulation remains. As more experimental evidence becomes available, these assumptions can be revisited to further improve the physiological fidelity of the model.

### 4.5. Conclusion and future work

Using a multi-scale physics-based computational model, we investigated how smooth muscle cell (SMC) mechanisms contribute to cerebral autoregulation across organizational scales, testing the hypothesis of Klein et al.^5^ that vessel-size-dependent pressure-diameter responses shape the organ-scale pressure-flow relationship and arise from differences in SMC abundance along the vascular tree. Our results support the first part of this hypothesis, as improved reproduction of the vessel-size-dependent pressure-diameter responses (Sections 3.1 and 4.1) consistently translated into more accurate organ-scale behavior (Sections 3.2 and 4.2). In contrast, the results do not support differences in SMC abundance as the main source of the observed vessel-size dependence, instead indicating a key role for functional heterogeneity in SMC behavior.

Beyond these specific hypotheses, the results demonstrate the value of mechanistic multi-scale modeling for linking cellular mechanisms to organ-scale hemodynamics, which is difficult to achieve experimentally due to scale separation. The sensitivity analysis further highlights the importance of a multi-scale perspective for understanding cerebral autoregulation by showing that the parameters with the strongest influence on autoregulatory behavior are distributed across all scales. To further expand the applicability of the model, future work should extend it from autoregulation to CBF regulation by incorporating additional regulatory pathways, such as endothelial and neurovascular control, and adapt it to represent pathophysiological conditions.

## 5. Author contribution

**Nele Demeersseman:** Methodology, Software, Formal analysis, Investigation, Validation, Data curation, Visualization, Writing – Original Draft, Writing – Review&Editing, Conceptualization, Funding acquisition; **Lauranne Maes:** Methodology, Writing – Review&Editing, Supervision; **Bart Depreitere:** Writing – Review&Editing, Conceptualization, Supervision; **Nele Famaey:** Writing – Review&Editing, Conceptualization, Supervision, Resources, Funding acquisition

## 8. Data availability statement

The MATLAB code of the multi-scale model will be made openly accessible upon journal acceptance. All other data required to reproduce the results are contained within the manuscript and supplementary information. If any clarification or additional information is needed, these can be obtained from the corresponding author upon request.

## 9. Supplementary information

One supplementary file is provided containing additional details on model formulation and calibration, along with supplementary results.

