## Supplementary material for "The Role of Smooth Muscle Cell Heterogeneity in Cerebral Autoregulation: A Multi-Scale Physics-Based Modeling Study"

### S1. Model formulation and implementation

#### S1.1. Organ scale

The cerebrovascular network geometry is generated using Murray's law and symmetric bifurcation, as described in the main text (Section 2.1.2.). Vessel lengths and wall thicknesses are assigned using segment-specific length-to-diameter and thickness-to-diameter ratios.

Apparent blood viscosity, $\mu$ [Pa·s], is calculated as a function of vessel diameter $D$ [$\mu$m], plasma viscosity $\mu_{plasma}$ [Pa·s], and discharge hematocrit $H_D$ using the *in vivo* empirical relations proposed by Pries et al.[1]:

$$\mu = \mu_{plasma}\left(1 + (w-1)\left(\frac{(1-H_D)^f - 1}{(1-0.45)^f - 1}\right)C\right)C \quad \text{(S1)}$$

with

$$f = \left(0.8 + e^{-0.075D^*}\right)\left(-1 + \frac{1}{1+10^{-11}D^{*12}}\right) + \frac{1}{1+10^{-11}D^{*12}}, \quad \text{(S2)}$$

$$w = 6e^{-0.085D^*} + 3.2 - 2.44e^{-0.06D^{*0.645}}, \quad \text{(S3)}$$

$$C = \left(\frac{D^*}{D^*-1.1}\right)^2, \quad \text{(S4)}$$

$$D^* = \frac{D}{1\ \mu m}, \quad \text{(S5)}$$

Assuming laminar flow, the hydraulic resistance of an individual vessel segment is calculated as:

$$R = \frac{8\mu L}{\pi r^4}, \quad \text{(S6)}$$

where $L$ is vessel length and $r$ is vessel radius.

Because each branch level consists of multiple parallel vessels with identical geometric properties, an effective resistance is computed for each branch level as:

$$R_{eff,j} = \frac{R_j}{N_j}, \quad \text{(S7)}$$

with $R_j$ the resistance of a single vessel and $N_j$ the number of vessels within branch level $j$.

Nodal pressures are determined from the prescribed inlet ($P_{in}$) and outlet ($P_{out}$) pressures by enforcing conservation of flow throughout the network:

$$P_k = \frac{R_{dist,k}P_{in} + R_{prox,k}P_{out}}{R_{prox,k} + R_{dist,k}}, \quad \text{(S8)}$$

where $R_{prox,k}$ and $R_{dist,k}$ denote the cumulative effective resistances between node k and the inlet and outlet boundaries, respectively.

Blood flow through each vessel is subsequently calculated using the Hagen-Poiseuille relation:

$$Q = \frac{\pi r^4}{8\mu L}(P_{prox} - P_{dist}), \tag{S9}$$

where $P_{prox}$ and $P_{dist}$ are the pressures at the proximal and distal ends of the vessel.

#### S1.2. Vessel and microstructural scale

For each vessel wall constituent $i$, the total deformation from its stress-free configuration to the current configuration is described by the deformation gradient

$$\boldsymbol{F}_i = \boldsymbol{F}_{vessel}\boldsymbol{G}_i, \tag{S10}$$

where $\boldsymbol{G}_i$ is the constituent-specific deposition stretch tensor that maps constituent $i$ from its stress-free configuration to the vessel reference configuration at baseline pressure, and $\boldsymbol{F}_{vessel}$ maps the vessel from this reference configuration to its current configuration under the applied pressure state $P_{l,network}$[2]. Assuming axisymmetric deformation without shear,

$$\boldsymbol{F}_{vessel} = \begin{bmatrix} \lambda_r & 0 & 0 \\ 0 & \lambda_\theta & 0 \\ 0 & 0 & \lambda_z \end{bmatrix}, \tag{S11}$$

where $\lambda_r$, $\lambda_\theta$, and $\lambda_z$ denote the radial, circumferential, and axial stretches, respectively[3]. The axial stretch is fixed as $\lambda_z = 1$, reflecting the assumption of constant vessel length under changing pressure conditions[4]. To account for through-wall variations in circumferential stretch, the vessel wall is discretized into 10 equally spaced radial points, at which the local stretch is calculated as $\lambda_\theta = \frac{r}{R}$, where $r$ and $R$ denote the local radial coordinates in the current and reference configurations, respectively[3]. Assuming incompressibility, the radial stretch is obtained as $\lambda_r = \frac{1}{\lambda_\theta \lambda_z}$[3].

For elastin, the deposition stretch tensor is defined as

$$\boldsymbol{G}_{elas} = \begin{bmatrix} g_{r,el} & 0 & 0 \\ 0 & g_{\theta,el} & 0 \\ 0 & 0 & g_{z,el} \end{bmatrix}, \tag{S12}$$

where $g_{r,el}$, $g_{\theta,el}$, and $g_{z,el}$ denote the radial, circumferential, and axial deposition stretches, respectively. The axial deposition stretch reflects axial *in vivo* pre-stretch and is assigned a fixed value from literature (see Table 1 in main text)[5]. The circumferential stretch $g_{\theta,el}$ is assumed constant over the vessel wall and obtained by solving the mechanical equilibrium equation in the baseline configuration, for which $\boldsymbol{F}_{vessel} = \mathbf{I}$[2]. The radial stretch is obtained as $g_{r,el} = \frac{1}{g_{\theta,el} g_{z,el}}$, assuming incompressiblity[3].

The Neo-Hookean strain-energy density function (Equation 6 in the main text) used to describe the constitutive behavior of elastin depends on the first invariant

$$I_{1,el} = \mathrm{trace}(\boldsymbol{C}_{el}), \tag{S13}$$

where

$$\boldsymbol{C}_{el} = \boldsymbol{F}_{el}{}^{T}\boldsymbol{F}_{el}\,, \tag{S14}$$

is the right Cauchy-Green deformation tensor of elastin[2].

Collagen is represented by two symmetrically oriented fiber families. For each fiber family $j$, the deposition stretch tensor is defined as

$$\boldsymbol{G}_{col,j} = g_{col}\boldsymbol{M}_{col,j} \otimes \boldsymbol{M}_{col,j} + \frac{1}{\sqrt{g_{col}}}\left(\boldsymbol{I} - \boldsymbol{M}_{col,j} \otimes \boldsymbol{M}_{col,j}\right), \tag{S15}$$

where

$$\boldsymbol{M}_{col,j} = [0 \quad \cos(\alpha_{col,j}) \quad \sin(\alpha_{col,j})]^{T}, \tag{S16}$$

represents the main fiber orientation in the reference configuration, specified by the fiber angle $\alpha_{col,j}$ relative to the circumferential direction. Assuming two symmetric fiber families, $\alpha_{col,2} = -\alpha_{col,1}$ [2].

The collagen strain-energy density function (Equation 7 in main text) depends on the first invariant

$$I_{1,col,j} = \mathrm{trace}\left(\boldsymbol{C}_{col,j}\right), \tag{S17}$$

and the fiber stretch invariant

$$I_{4,col,j} = \boldsymbol{M}_{col,j}{}^{T}\left(\boldsymbol{C}_{col,j}\boldsymbol{M}_{col,j}\right), \tag{S18}$$

where

$$\boldsymbol{C}_{col,j} = \boldsymbol{F}_{col,j}{}^{T}\boldsymbol{F}_{col,j}, \tag{S19}$$

is the right Cauchy-Green deformation tensor of collagen[2].

The smooth muscle cells (SMCs) are assumed to be deposited in the reference configuration and only feel deformation with respect to that configuration[2] such that

$$\boldsymbol{G}_{SMC,j} = \boldsymbol{I}, \tag{S20}$$

and thus

$$\boldsymbol{F}_{SMC} = \boldsymbol{F}_{vessel}. \tag{S21}$$

The SMC strain-energy density function (Equation 8 in main text) depends on the invariant

$$I_{4,SMC,j} = \boldsymbol{M}_{SMC,j}{}^{T}\left(\boldsymbol{C}_{col}\boldsymbol{M}_{SMC,j}\right), \tag{S22}$$

where

$$\boldsymbol{C}_{SMC} = \boldsymbol{F}_{SMC}{}^{T}\boldsymbol{F}_{SMC}, \tag{S23}$$

is the right Cauchy-Green deformation tensor of the SMC constituent and

$$\boldsymbol{M}_{SMC,j} = [0 \quad \cos(\alpha_{SMC,j}) \quad \sin(\alpha_{SMC,j})]^T, \tag{S24}$$

denotes the orientation of SMC family $j$ with $\alpha_{SMC,j}$ the angle relative to the circumferential direction[2]. Assuming two symmetric SMC families, $\alpha_{SMC,2} = -\alpha_{SMC,1}$.

The normalized relative sliding between SMC filaments, $u_{rs,j}$, is determined from the steady-state balance between the passive material resistance,

$$P_{mat,j} = \mu_{SMC}(n_3 + n_4)(\sqrt{I_{4,SMC,j}} + \mu_{rs,j} - 1), \tag{S25}$$

and the active cross-bridge contribution,

$$P_{SMC,j} = \begin{cases} \kappa_{SMC} n_3 & \text{if } P_{mat,j} < \kappa_{SMC} n_3 \\ P_{mat,j} & \text{if } \kappa_{SMC} n_3 \leq P_{mat,j} \leq \kappa_{SMC}(n_3 + n_4), \\ \kappa_{SMC}(n_3 + n_4) & \text{if } P_{mat,j} > \kappa_{SMC}(n_3 + n_4) \end{cases} \tag{S26}$$

where $\kappa_{SMC}$ is a parameter characterizing the driving force of individual cross-bridges[2,6]. When $P_{mat,j} < \kappa_{SMC} n_3$, the steady-state value of $u_{rs,j}$ is

$$\mu_{rs,j} = \frac{\kappa_{SMC} n_3}{\mu_{SMC}(n_3+n_4)} + 1 - \sqrt{I_{4,SMC,j}}, \tag{S27}$$

such that $P_{mat,j} = \kappa_{SMC} n_3$ and equilibrium ($P_{SMC,j} = P_{mat,j}$) is obtained. Similarly, when $P_{mat,j} > \kappa_{SMC}(n_3 + n_4)$,

$$\mu_{rs,j} = \frac{\kappa_{SMC}}{\mu_{SMC}} + 1 - \sqrt{I_{4,SMC,j}}, \tag{S28}$$

such that $P_{mat,j} = \kappa_{SMC}(n_3 + n_4) = P_{SMC,j}$. For intermediate values $\kappa_{SMC} n_3 \leq P_{mat,j} < \kappa_{SMC}(n_3 + n_4)$, the equilibrium condition is already satisfied and $u_{rs,j}$ remains unchanged[2].

#### S1.3. Molecular scale

Cross-bridge cycling in smooth muscle cells (SMCs) is described using the four-state cross-bridge model of Hai and Murphy, consisting of detached dephosphorylated cross-bridges ($n_1$), detached phosphorylated cross-bridges ($n_2$), attached phosphorylated cross-bridges ($n_3$), and attached dephosphorylated cross-bridges ($n_4$)[7]. The dynamics of the four states are governed by:

$$\frac{d}{dt}\begin{bmatrix} n_1 \\ n_2 \\ n_3 \\ n_4 \end{bmatrix} = \begin{bmatrix} -k_1 & k_2 & 0 & k_7 \\ k_1 & -(k_2 + k_3) & k_4 & 0 \\ 0 & k_3 & -(k_4 + k_5) & k_6 \\ 0 & 0 & k_5 & -(k_6 + k_7) \end{bmatrix}\begin{bmatrix} n_1 \\ n_2 \\ n_3 \\ n_4 \end{bmatrix}, \tag{S29}$$

where $k_1$ to $k_7$ are rate constants. The state variables represent fractions of the total cross-bridge population and therefore satisfy:

$$n_1 + n_2 + n_3 + n_4 = 1. \tag{S30}$$

The phosphorylation rate $k_1$ depends on intracellular calcium concentration as described in the main text (Equations 9 and 10). For the simulation scenario incorporating functional SMC heterogeneity, this calcium concentration is calculated by combining Equations 12 and 13 of the main text as:

$$[Ca^{2+}] = [Ca^{2+}_{base}(D)] \cdot A_{CaP} \frac{{P_{T,n}}^{n_{CaP}(D,P_{T,n})}}{{P_{T,n}}^{n_{CaP}(D,P_{T,n})} + {K_{CaP}}^{n_{CaP}(D,P_{T,n})}}. \tag{S31}$$

Vessel-size-dependency of $n_{CaP}$ and $[Ca^{2+}_{base}]$ is inferred by interpolation between experimental measurements obtained for one small- and one large-diameter vessel size[8,9]. Because data are only available for these two vessel sizes, a shape parameter $s$ is introduced to control the transition between the small- and large-diameter data, such that:

$$n_{CaP}(D, P_{T,n}) = \begin{cases} n_{CaP,L} & \text{if } P_{T,n} \leq 1 \\ \max\left(0, n_{CaP,H,s} + (n_{CaP,H,l} - n_{CaP,H,s}) \frac{D^{s_N} - {D_{CaP,s}}^{s_N}}{{D_{CaP,l}}^{s_N} - {D_{CaP,s}}^{s_N}}\right) & \text{if } P_{T,n} > 1 \end{cases} \tag{S32}$$

$$[Ca^{2+}_{base}(D)] = [Ca_{base,s}] + ([Ca_{base,l}] - [Ca_{base,s}]) \frac{D^{s_{Ca}} - {D_{CaP,s}}^{s_{Ca}}}{{D_{CaP,l}}^{s_{Ca}} - {D_{CaP,s}}^{s_{Ca}}}, \tag{S33}$$

where $D$ is vessel diameter and the subscripts $s$ and $l$ denote parameter values associated with the experimental small- and large-diameter vessels, respectively.

#### S1.4. Convergence criteria

For each prescribed inlet pressure, the multi-scale problem is solved iteratively. During each iteration, vessel diameters are updated in response to local pressure conditions. The updated diameters are then fed back to the organ-scale vascular network, resulting in a redistribution of vascular resistances and pressures. This pressure-diameter coupling is repeated until convergence of the vessel diameters is achieved. Convergence is assessed based on the mean relative change in vessel diameter between two successive iterations:

$$\frac{1}{N} \sum_{i=1}^{N} \left| \frac{D_i^{(k)} - D_i^{(k-1)}}{D_i^{(k-1)}} \right| \leq \varepsilon, \tag{S34}$$

where $D_i^{(n)}$ is the diameter of vessel $i$ at iteration $k$, $N$ is the total number of vessels in the network, and $\varepsilon$=$10^{-3}$ is the convergence tolerance. To prevent indefinite iterations for non-convergent parameter combinations encountered during the sensitivity analysis, a maximum of 100 iterations is imposed. Simulations reaching this limit are terminated and flagged for further inspection.

### S2. Parameter calibration and sensitivity analysis

The following paragraphs provide additional justification for some parameter values and sensitivity ranges assigned in Table 1 of the main text.

The number of capillaries per terminal arteriole ($N_c$) is not derived from a single anatomical measurement. Instead, it is calibrated such that the model reproduces physiological pressure[10] and wall shear stress[11–13] distributions, as commonly done in vascular models[14,15].

To ensure physically consistent sensitivity analysis of vessel wall composition, the elastin ($\phi_{el}$), collagen ($\phi_{col}$), and smooth muscle cell ($\phi_{SMC}$) volume fractions are reparametrized, as independent variation would violate their constraint of summing to unity ($\phi_{el} + \phi_{col} + \phi_{SMC} = 1$). $\phi_{SMC}$ and the fraction $\theta_{e/c} = \frac{\phi_{el}}{\phi_{el}+\phi_{col}}$ are treated as independent parameters, from which $\phi_{el}$ and $\phi_{col}$ are determined as $(1 - \phi_{SMC})\theta_{e/c}$ and $(1 - \phi_{SMC})(1 - \theta_{e/c})$, respectively.

The parameters $CaM_{max}$ and $K_{CaM}$ are not uniquely established for cerebrovascular SMCs, nor for the loading and activation conditions considered in this study. However, Sobue et al.[16] suggested that their values are likely to fall within the ranges of 2-10 µM and 0.5-5 µM, respectively. To obtain physiologically realistic calcium-dependent phosphorylation behavior across simulation scenarios, $CaM_{max}$ and $K_{CaM}$ are optimized within these ranges for each scenario. Specifically, a joint optimization is performed to (i) align the point of maximum sensitivity (maximal slope) of the calcium-$k_1$ relationship (Equations 9 and 10 in main text) with the median baseline intracellular calcium concentration across the vascular network and (ii) obtain a phosphorylation rate of approximately 0.2 $s^{-1}$ ($= k_{1,base}$) at this calcium concentration[6]. The first criterion reflects the expectation that cerebrovascular SMCs operate most responsively around their baseline intracellular calcium concentration. To render the optimization well posed, $k_{1,max}$ is fixed to 0.5 $s^{-1}$. During sensitivity analysis, uncertainty in $CaM_{max}$ and $K_{CaM}$ is explored by varying $k_{1,base}$ and calculating the corresponding $CaM_{max}$ and $K_{CaM}$.

For the functional SMC heterogeneity scenario, the pressure-calcium relationship (Equation 12 in main text) is expressed in normalized coordinates and therefore constrained to pass through the baseline operating point ($P_{T,n}, Ca_n^{2+}$)=(1,1). This constraint allows $A_{CaP}$ to be written as

$$A_{CaP} = 1 + {K_{CaP}}^{n_{CaP}(D,P_{T,n})}, \tag{S35}$$

such that only $K_{CaP}$ and $n_{CaP}$ still need to be estimated during curve fitting[9].

As described in section S1.3. Molecular scale, the diameter dependencies of $n_{CaP}$ and $[Ca_{base}^{2+}]$ are obtained by interpolating between experimental measurements from one small- and one large-diameter vessel size[8,9]. Because there is only data for two vessel sizes, the shape parameters in Equations S32 and S33 cannot be uniquely defined. During calibration, the shape parameters are selected such that the resulting interpolations correspond to parabolic transitions between the small- and large-vessel data. During sensitivity analysis, uncertainty in the shape of the diameter-dependent functions is explored by varying the shape parameters over a symmetric range around the linear case ($s = 1$). The remaining parameters defining Equations S32 and S33 are associated with the experimental small- and large-vessel datasets. As these datasets were obtained in rats[9], species-specific differences are accounted for where possible during calibration, consistent with the aim of developing a human-relevant model (see Section 2.3. in the main text). The large-vessel measurements were acquired in rat middle cerebral arteries (MCAs)[9], and therefore the corresponding reference diameter $D_{CaP,l}$ is taken equal to the diameter of the human MCA[17]. The small-vessel dataset was obtained in rat parenchymal arterioles with a diameter of 36 µm[9]. Assuming the same scaling laws as used for capillary diameters, this translates to a diameter of 63 µm in humans[18] and 50 µm in piglets[18,19].

### S3. Alternative implementations of the homogeneous SMC scenario

As described in Section 2.2. in the main text, four experimental pressure-calcium relationships reported by Cipolla et al.[8] and Nystoriak et al.[9] can be used to describe the homogeneous SMC scenario. The main manuscript uses the arteriole data of Nystoriak et al.[9]. Here results are generated for all four relationships to assess the sensitivity of the homogeneous SMC scenario to the selected experimental dataset. Figure 1 shows all relationships and their corresponding fit.

The pressure-calcium ($P_T - Ca^{2+}$) relationship obtained from the Nystoriak arteriole data[9] is fitted by

$$[Ca^{2+}] = A_{CaP}^{hom} \frac{{P_T}^{n_{CaP}^{hom}}}{{P_T}^{n_{CaP}^{hom}} + {K_{CaP}^{hom}}^{n_{CaP}^{hom}}}, \tag{S36}$$

with $A_{CaP}^{hom}$ = 359.01 nM, $K_{CaP}^{hom}$ = 13.37 mmHg, and $n_{CaP}^{hom}$ = 0.78, as shown in the main text.

The middle cerebral artery data of Nystoriak et al.[9] are fitted by

$$[Ca^{2+}] = aP_T + b, \tag{S37}$$

with $a$ = 1.35 nM/mmHg and $b$ = 111.7 nM.

The arteriole data of Cipolla et al.[8] are fitted by

$$[Ca^{2+}] = A_{CaP} \frac{{P_T}^{n_{CaP}}}{{P_T}^{n_{CaP}} + {K_{CaP}}^{n_{CaP}}}, \tag{S38}$$

with $A_{CaP}$ = 290.95 nM, $K_{CaP}$ = 20.72 mmHg, and $n_{CaP}$ = 0.81.

Finally, the middle cerebral artery data of Cipolla et al.[8] are fitted by

$$[Ca^{2+}] = a + A_{CaP} \frac{{P_T}^{n_{CaP}}}{{P_T}^{n_{CaP}} + {K_{CaP}}^{n_{CaP}}}, \tag{S39}$$

with $a$ = 92.94 nM, $A_{CaP}$ = 192.91 nM, $K_{CaP}$ = 112.72 mmHg, and $n_{CaP}$ = 3.48.

Figure 2 shows the vessel-scale pressure-diameter responses for the homogeneous SMC scenario using the four possible experimental pressure-calcium relationships of Figure 1. Although the selected relationship affects the diameter responses quantitatively, it does not alter the conclusions drawn in the main text. Across all four parameterizations, homogeneous SMC activation introduces active diameter regulation, with dilation at reduced pressures and constriction at elevated pressures, while failing to reproduce the experimentally observed vessel-size-dependent trends[20]. Introducing structural SMC heterogeneity (Figure 3) consistently reduces both dilation at reduced pressures and constriction at elevated pressures in smaller vessels. This improves agreement with the experimental size-dependent trends at elevated pressures but worsens it at reduced pressures. Thus, the main-text conclusion that functional SMC heterogeneity is required to reproduce the experimentally observed vessel-size-dependent trends in pressure-diameter responses holds irrespective of the pressure-calcium relationship selected for the homogeneous scenario.

The similar qualitative behavior across the four parameterizations can be understood from the vessel-specific baseline calcium levels and pressure sensitivities (pressure-calcium slopes) they impose. As discussed for the functional SMC heterogeneity scenario in Section 4.1. of the main text, these features determine how pressure changes are translated into SMC activation. In contrast to the functional heterogeneity scenario, all four homogeneous parameterizations assign lower baseline calcium levels to smaller vessels and do not reverse which vessel size class exhibits the greatest pressure sensitivity across the pressure range. This results in the same qualitative vessel-size-dependent trends across parameterizations, while differences in the exact baseline calcium levels and pressure sensitivities account for quantitative differences in the diameter responses. For example, the Cipolla[8] artery-based parameterization is the only one in which the pressure-calcium slope increases with increasing pressure (Figure 1), and correspondingly the only one showing a stronger diameter response to pressure elevations than to pressure reductions (Figure 2).

At the organ-scale, as at the vessel-scale, the choice of pressure-calcium parameterization affects the pressure-diameter and pressure-flow responses quantitatively, but does not alter the overall conclusions. The homogeneous SMC scenario provides better agreement with the experimental responses[20] than the passive SMC scenario and poorer agreement than the functional SMC heterogeneity scenario, as reflected by the $R^2$ and nRMSE metrics (Figure 4). The correspondence between vessel- and organ-scale responses is also maintained across all four parametrizations. This is particularly evident for structural SMC heterogeneity, for which the improved agreement at elevated pressures and poorer agreement at reduced pressures observed at the vessel-scale are reflected in the organ-scale responses (Figure 5).

### S4. Sensitivity analysis

This section provides a more detailed view of the pressure dependence of parameter sensitivity. For different levels of cerebral perfusion pressure, the top ten parameters according to the mean of the absolute elementary effects ($\mu^*$), the standard deviation of the elementary effects ($\sigma$), and their geometric mean after normalization ($\sqrt{\mu_n^* \sigma_n}$), are provided in Figure 6 panels A, B, and C, respectively. Comparison of panels A and B shows substantial overlap among the highest-ranking parameters, indicating that parameters with a strong overall influence on model output ($\mu^*$) often also exhibit pronounced nonlinearity or interaction with other parameters ($\sigma$). As discussed in Section 4.3 of the main text, panel C shows that organ-scale parameters become increasingly represented among the highest-ranking parameters as pressure decreases, while molecular-scale parameters become less prominent.

### Figures

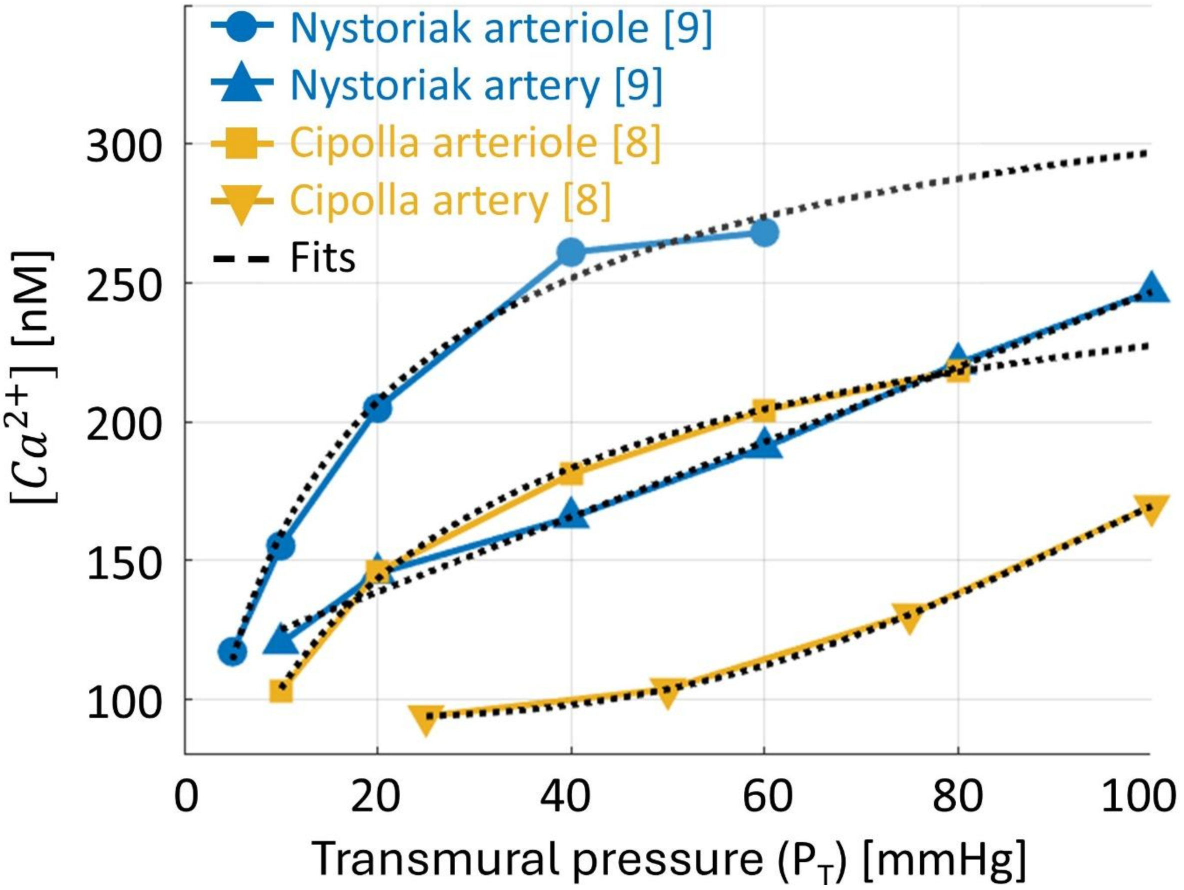


***Figure 1*** *Experimental pressure-calcium relationships Experimental pressure-calcium relationships reported by Cipolla et al.*[8] *and Nystoriak et al.*[9] *for arterioles and middle cerebral arteries, together with their corresponding fitted functions (Equations S36-S39). Nystoriak et al.*[9] *data are shown as blue circles (parenchymal arterioles) and blue upward triangles (middle cerebral arteries), while Cipolla et al.*[8] *data are shown as yellow squares (parenchymal arterioles) and yellow downward triangles (middle cerebral arteries). Black dotted lines indicate the corresponding fitted functions. The four datasets are used to assess the sensitivity of the homogeneous SMC scenario to the selected experimental pressure-calcium relationship.*

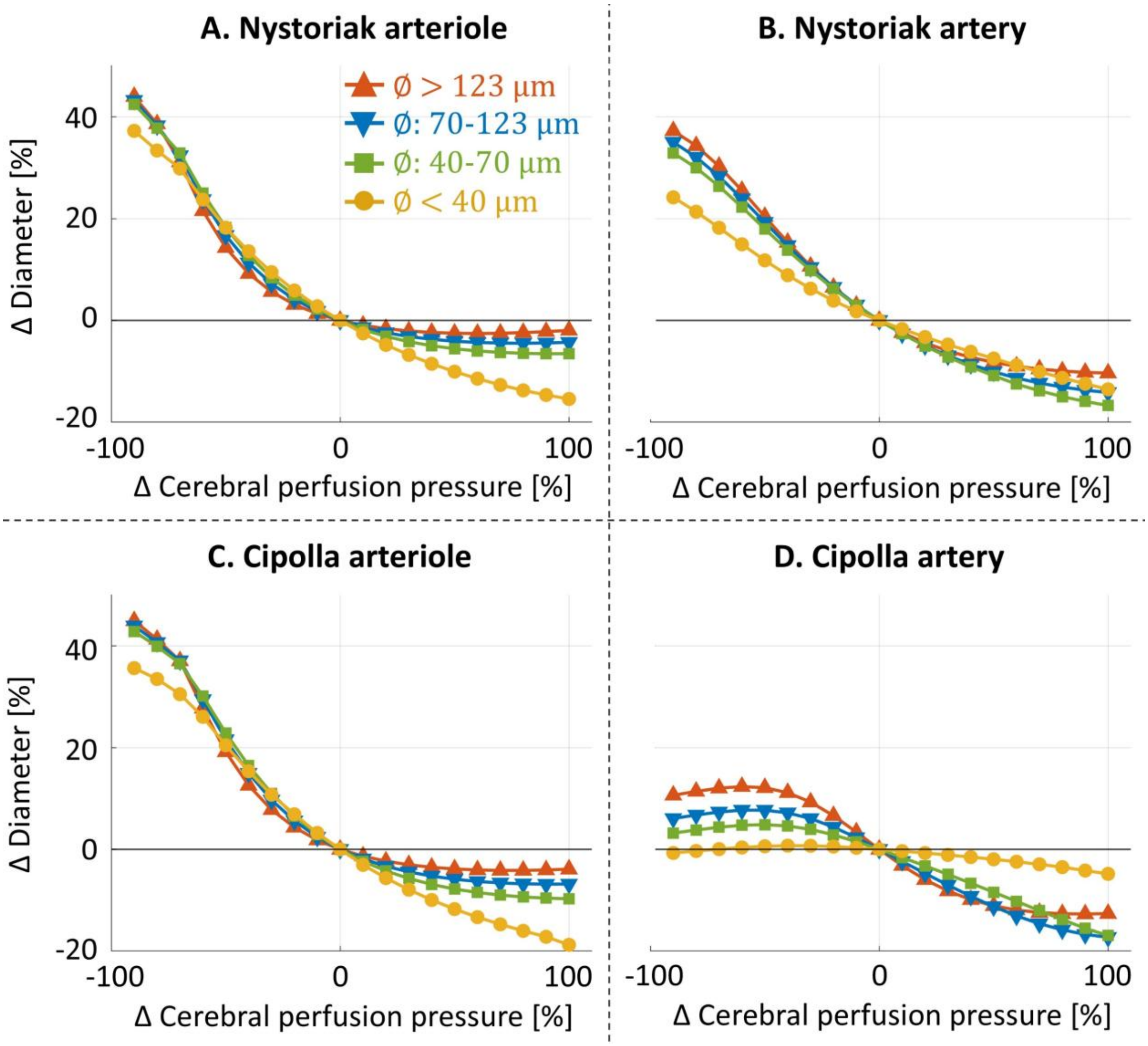


***Figure 2*** *Vessel diameter (relative to baseline) as a function of cerebral perfusion pressure (relative to baseline) for the homogeneous SMC scenario using four alternative experimental pressure-calcium relationships. Panels A to D correspond to the results obtained using pressure-calcium relationships fitted to the parenchymal arteriole data of Nystoriak et al.*[9]*, the middle cerebral artery data of Nystoriak et al.* [9]*, the parenchymal arteriole data of Cipolla et al.*[8]*, and the middle cerebral artery data of Cipolla et al.* [8]*, respectively. Within each panel, the different curves represent the average pressure-diameter response of all vessels within a given size class: diameters <40 µm in yellow circles, 40-70 µm in green squares, 70-123 µm in blue downward triangles, and >123 µm in red upward triangles.*

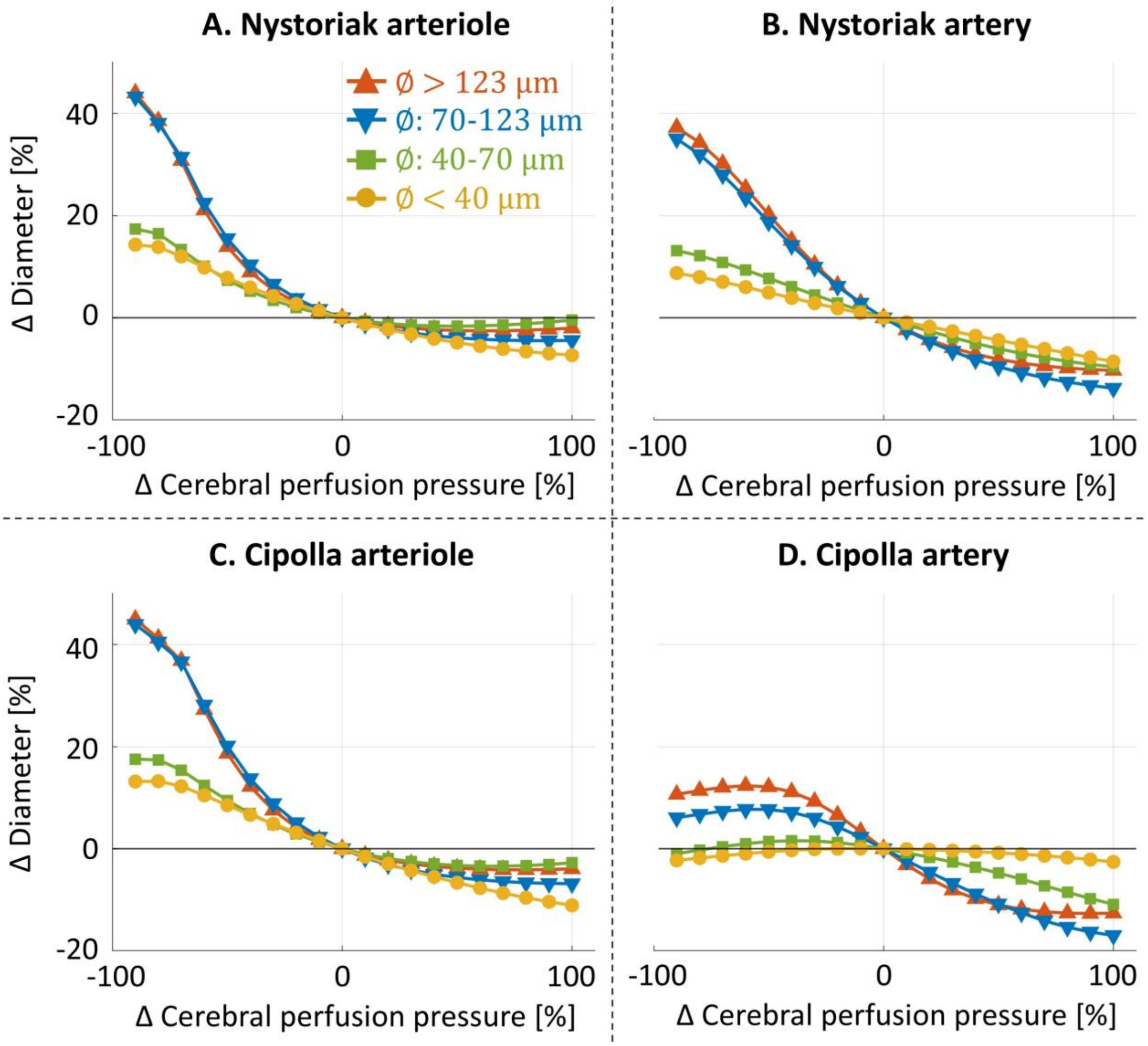


***Figure 3*** *Vessel diameter (relative to baseline) as a function of cerebral perfusion pressure (relative to baseline) for the structural SMC heterogeneity scenario using four alternative experimental pressure-calcium relationships. Panels A to D correspond to the results obtained using pressure-calcium relationships fitted to the parenchymal arteriole data of Nystoriak et al.[9], the middle cerebral artery data of Nystoriak et al. [9], the parenchymal arteriole data of Cipolla et al.[8], and the middle cerebral artery data of Cipolla et al. [8], respectively. Within each panel, the different curves represent the average pressure-diameter response of all vessels within a given size class: diameters <40 μm in yellow circles, 40-70 μm in green squares, 70-123 μm in blue downward triangles, and >123 μm in red upward triangles.*

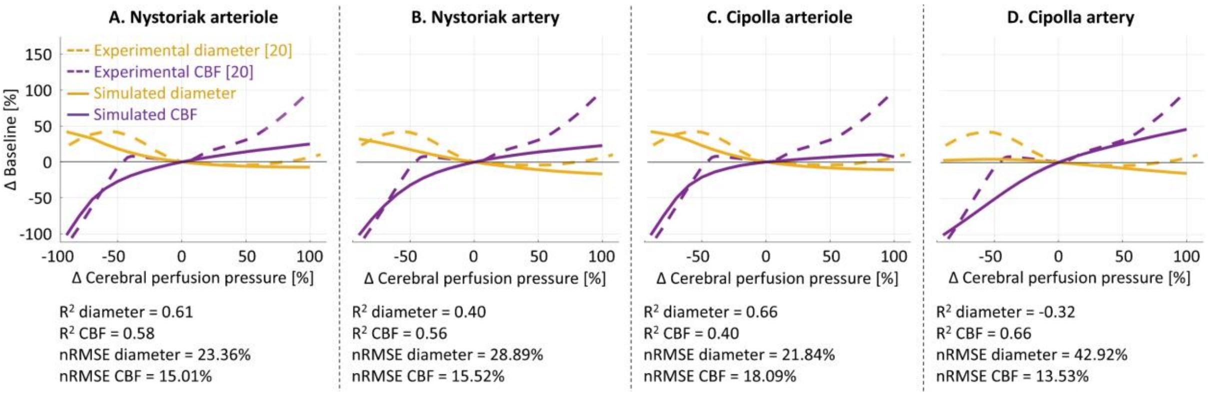


***Figure 4*** *Organ-scale pressure-flow and pressure-diameter curves (relative to baseline) for the homogeneous SMC scenario using four alternative experimental pressure-calcium relationships. Model predictions (full lines) are compared to experimental data[20] (dashed lines), with cerebral blood flow (CBF) shown in purple and vessel diameter in yellow. Panels A to D correspond to the results obtained using pressure-calcium relationships fitted to the parenchymal arteriole data of Nystoriak et al.[9], the middle cerebral artery data of Nystoriak et al. [9], the parenchymal arteriole data of Cipolla et al.[88], and the middle cerebral artery data of Cipolla et al. , respectively. Quantitative agreement between model and experiments is reported using the coefficient of determination (R²) and the normalized root mean square error (nRMSE).*

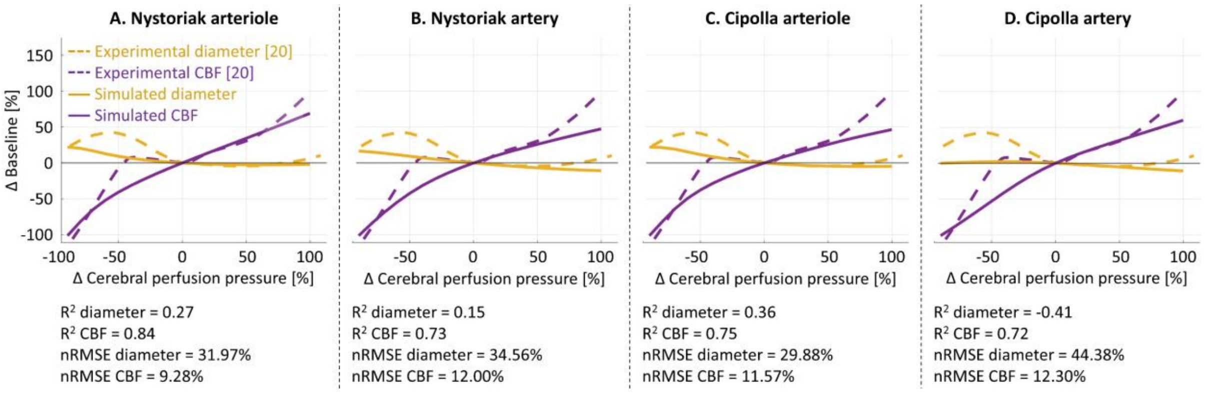


***Figure 5*** *Organ-scale pressure-flow and pressure-diameter curves (relative to baseline) for the structural SMC heterogeneity scenario using four alternative experimental pressure-calcium relationships. Model predictions (full lines) are compared to experimental data[20] (dashed lines), with cerebral blood flow (CBF) shown in purple and vessel diameter in yellow. Panels A to D correspond to the results obtained using pressure-calcium relationships fitted to the parenchymal arteriole data of Nystoriak et al.[9], the middle cerebral artery data of Nystoriak et al. [9], the parenchymal arteriole data of Cipolla et al.[8], and the middle cerebral artery data of Cipolla et al. [8], respectively. Quantitative agreement between model and experiments is reported using the coefficient of determination (R²) and the normalized root mean square error (nRMSE).*

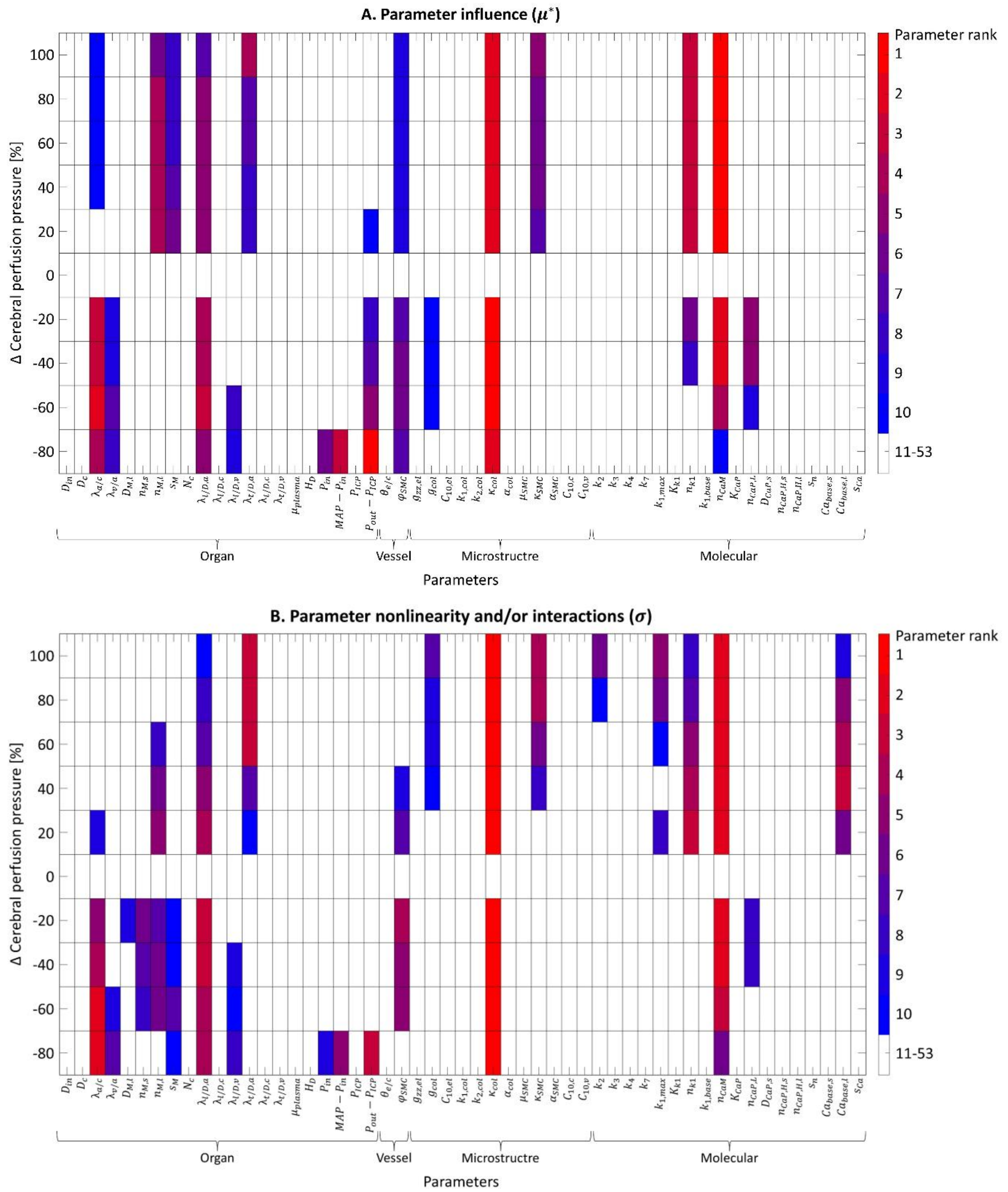


***Figure 6*** (Part 1) Pressure-dependent parameter sensitivity for the functional SMC heterogeneity scenario. Parameter rankings at different levels of cerebral perfusion pressure based on (A) the mean of the absolute elementary effects ($\mu^*$) and (B) the standard deviation of the elementary effects ($\sigma$). For each pressure level, the ten parameters with the highest value of the respective sensitivity measure are highlighted according to the color coding shown on the right. Parameters are grouped according to the model scale they relate to: organ, vessel, microstructure, and molecular.

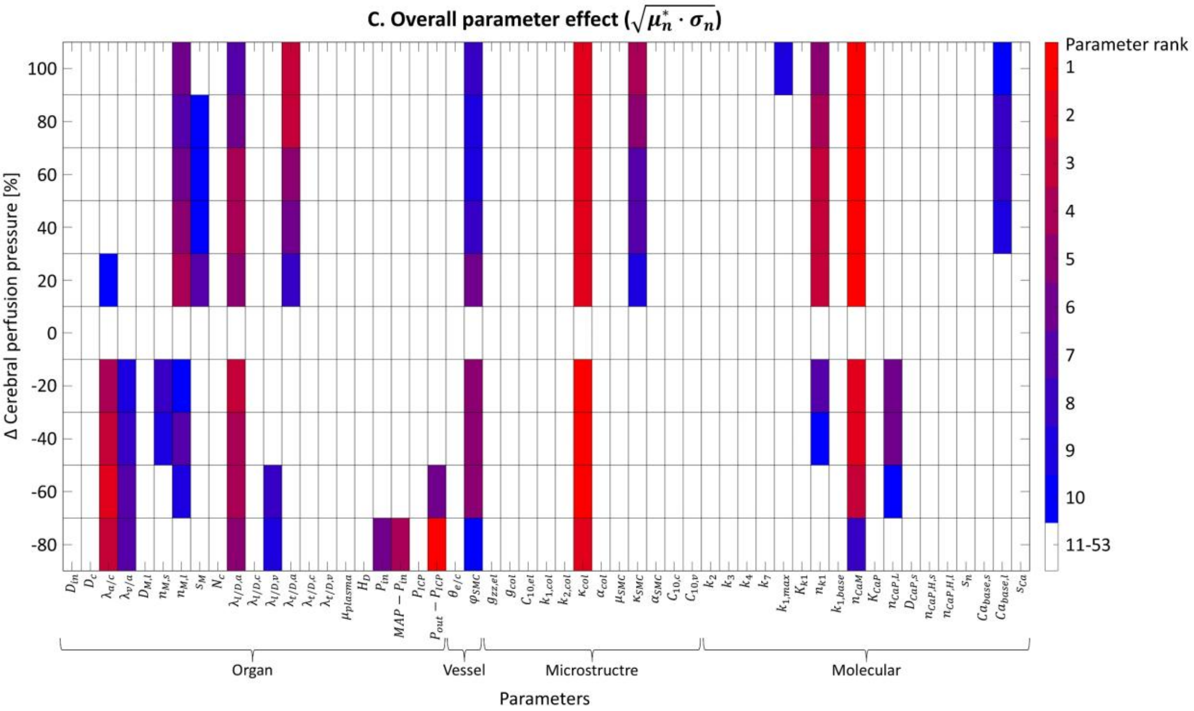


*Figure 6* (Part 2) Pressure-dependent parameter sensitivity for the functional SMC heterogeneity scenario. Parameter rankings at different levels of cerebral perfusion pressure based on (C) the geometric mean of the normalized mean of the absolute elementary effects μ* and the normalized standard deviation of the elementary effects σ. For each pressure level, the ten parameters with the highest value of the respective sensitivity measure are highlighted according to the color coding shown on the right. Parameters are grouped according to the model scale they relate to: organ, vessel, microstructure, and molecular.
